# Non-genetic maternal effects shape individual differences in cortisol phenotypes in wild chimpanzees

**DOI:** 10.1101/2021.07.16.452609

**Authors:** Patrick J. Tkaczynski, Fabrizio Mafessoni, Cédric Girard-Buttoz, Liran Samuni, Corinne Y. Ackermann, Pawel Fedurek, Cristina Gomes, Catherine Hobaiter, Therese Löhrich, Virgile Manin, Anna Preis, Prince D. Valé, Erin G. Wessling, Livia Wittiger, Zinta Zommers, Klaus Zuberbuehler, Linda Vigilant, Tobias Deschner, Roman M. Wittig, Catherine Crockford

## Abstract

Glucocorticoids, such as cortisol, mediate homeostatic processes, allowing individuals to adjust to fluctuating environments. The regulation of circadian cortisol responses, a key homeostatic function, has been shown to be heritable. However, to understand better the role of parental care in shaping physiological functioning in long-lived mammals with protracted parental care, there is a need to disentangle genetic and non-genetic parental contributions to variation in glucocorticoid phenotypes. We used a dataset of 6,123 cortisol measures from urine samples from 170 wild chimpanzees spanning 18 years of data collection. We found consistent inter-individual differences in circadian cortisol phenotypes, with differences most apparent when considering average cortisol levels given the effect of time of day. Maternal effects explained around 10% (2-18%) variation in these average cortisol levels, while variation attributable to genetic factors was not distinguishable from zero. Our results indicate, relative to genetic effects, a qualitatively stronger influence of mothers, whether via epigenetic processes or via behavioral priming for coping with stressors, in shaping cortisol phenotypes in this species. This provides novel insight into the vital role of mothers in the developmental plasticity of long-lived mammals and, more generally, the selective pressures shaping physiological plasticity.

## Introduction

In vertebrates, glucocorticoids (GCs), secreted via the hypothalamic-pituitary-adrenal (HPA) axis, facilitate homeostasis via mediation of metabolic, immune, and behavioral responses to intrinsic and extrinsic stressors (Sapolsky et al., 2000; Selye, 1976; Smith and Vale, 2006; Tsigos and Chrousos, 2002). As a consequence of this multi-faceted and dynamic role, the regulation of HPA axis activation and GC secretion is of broad interest to ecologists and evolutionary biologists seeking to understand how animals adapt to changing environments (Beehner and Bergman, 2017; Bonier and Cox, 2020; Bonier and Martin, 2016; Guindre-Parker, 2020, 2018; Guindre-Parker et al., 2019). Despite the flexibility of HPA axis activity in response to external and internal stimuli, numerous studies demonstrate consistent individual differences in HPA axis activity and reactivity to environmental stimuli (Schoenemann and Bonier, 2018; Taff et al., 2018). Recent evidence suggests that inter-individual variation in HPA axis regulation can be predictive of variation in fitness outcomes (Bonier and Cox, 2020; Campos et al., 2021). For example, female baboons with consistently elevated HPA axis activity live substantially shorter lives than those with lower HPA axis activity (Campos et al., 2021). Given the profound fitness effects of individual differences in HPA axis activity and regulation, understanding the relative role of genetics, experience, and environment in shaping these GC phenotypes is key to understanding the evolution of physiological plasticity (Bonier and Martin, 2016; Guindre-Parker, 2018).

In many wild animal populations, environmental heterogeneity can increase within-individual variation in average GC levels and mask between-individual differences (Baugh et al., 2014; Cook et al., 2012; Grace and Anderson, 2014; Montiglio et al., 2015; Sparkman et al., 2014; Taff et al., 2018; Tkaczynski et al., 2019). As a consequence, more recent studies have begun to focus on the degree in which individuals vary in GC secretion in response to shifting environmental gradients, i.e. GC reaction norms (Araya‐Ajoy et al., 2015; Araya-Ajoy and Dingemanse, 2017; Guindre-Parker, 2020; Guindre-Parker et al., 2019; Sonnweber et al., 2018). In humans, the circadian cortisol (the main GC in vertebrates) pattern is a well described reaction norm: levels rise gradually during sleep prior to a peak upon awakening, followed by declines throughout the day (Weitzman et al., 1971). A wealth of human studies reveal that deviations from this pattern, typically caused by a lack of a decline in cortisol levels during the latter half of the day, are related to poor physical and/or mental health (Butler et al., 2017; Carrion et al., 2002; Corbett et al., 2006; Gonzalez et al., 2009; Gustafsson et al., 2010; Saridjan et al., 2010; Sephton et al., 2000; Zilioli et al., 2016), and may also be predictive of survival (Sephton et al., 2000). Results from human twin studies indicate as much as 60% of the variation in circadian cortisol reactivity may be explained by genetic effects (Bartels et al., 2003a, 2003b; Gustafsson et al., 2011; Steptoe et al., 2009). While twin studies in humans have been important in revealing the genetic regulation of circadian cortisol responses, these studies are constrained in their ability to disentangle the genetic and non-genetic parental effects shaping this GC phenotype (Morris et al., 2020).

Circadian cortisol responses have recently begun to receive attention within non-human animal ecology (Behringer et al., 2020; Emery Thompson et al., 2020; Girard-Buttoz et al., 2021; Sonnweber et al., 2018). Species with protracted development phases and prolonged parental dependencies offer exciting opportunities to better quantify the relative influence of genetic or non-parental effects on circadian cortisol regulation. These insights can help us understand whether protracted development as a life history adaptation has led to, and potentially been selected for, a greater influence of parental effects on offspring physiology.

Parental, and in particular maternal, effects are recognized as major evolutionary drivers of trait variation (Moore et al., 2019). In experimental rodent studies, maternal cortisol levels during pregnancy and during post-partum offspring rearing, as well as rates of maternal interaction with offspring, are all predictors of offspring cortisol levels and reactivity (Champagne and Curley, 2009; Maccari et al., 2014). Rodent studies also suggest that maternal effects may occur via epigenetic processes, such as DNA methylation of GC receptor promotor regions, leading to altered responsivity to stressors (Champagne, 2008; Champagne and Curley, 2009; Zhang et al., 2013). Non-human primate (hereafter primate) studies of the role of maternal effects on cortisol secretion and reactivity have typically employed maternal deprivation paradigms, either via experimental separations or due to naturally occurring maternal loss (Champagne and Curley, 2009; Girard-Buttoz et al., 2020; Rosenbaum et al., 2020). Here, maternal loss is linked to elevations in cortisol levels or alterations to diurnal rhythm (Girard-Buttoz et al., 2020; Shannon et al., 1998), however, these effects do not necessarily last into adulthood (Girard-Buttoz et al., 2020; Rosenbaum et al., 2020). Similarly, in human studies, tests of maternal effects on cortisol regulation classically examine the consequences of negative maternal or early life circumstances (e.g. poor mental or physical health, low socioeconomic status, or maternal loss (reviewed in Champagne and Curley, 2009). Here, maternal loss or early life adversity related to maternal condition are associated with elevated HPA activity in offspring, which can last into adulthood for some individuals. Therefore, much of what we know about maternal, rather than genetic, effects on cortisol regulation in long-lived mammals is derived from studies of manipulated and/or extreme maternal circumstances.

In our study, we tackle the challenge of disentangling the relative contributions of genetic and non-genetic maternal effects to variation in cortisol phenotypes in wild chimpanzees. Like humans, chimpanzees are long-lived mammals with protracted developmental phases (Bründl et al., 2021; Crockford et al., 2020; Nakamura et al., 2014; Samuni et al., 2020; Stanton et al., 2020). In addition, many of the environmental factors influencing variation in cortisol levels in chimpanzees are established (Emery Thompson et al., 2020, 2010; Muller and Wrangham, 2004; Preis et al., 2019; Samuni et al., 2019; Sonnweber et al., 2018; Wessling et al., 2018a, 2018b), and, therefore, can be accounted and controlled for when modeling individual variation in cortisol phenotypes.

First, we examine whether there are consistent individual differences in circadian cortisol responses in five different communities and two subspecies of wild chimpanzees (western, *Pan troglodytes verus* and eastern, *Pan troglodytes schweinfurthii*). The dataset includes 170 individuals representing adults and immature individuals of both sexes. Using Bayesian analyses and permutation tests within the framework of an animal model approach (Wilson et al., 2010), we present estimates of the relative contributions of genetic, maternal, and environmental effects to circadian cortisol responses in this wild, long-lived mammal.

In chimpanzees, as in humans, cortisol secretion peaks with the awakening response, followed by a decline throughout the day (Muller and Lipson, 2003). Consistent individual differences in circadian cortisol responses are discernible in adult males (Sonnweber et al., 2018), and in both sexes, these patterns vary due to aging (Emery Thompson et al., 2020) during ill health (Behringer et al., 2020), or following traumatic events such as maternal loss during immaturity (Girard-Buttoz et al., 2021). Chimpanzees are a relatively long-lived species, have a gestation period of approximately 8 months, and a prolonged immature dependency lasting at least 10 years, in which there is emerging evidence of maternal influences in growth, survival, and future reproductive success (Crockford et al., 2020; Nakamura et al., 2014; Samuni et al., 2020; Stanton et al., 2020). Therefore, during both pre- and post-natal phases, there is a long period in which maternal and environmental factors can shape endocrine phenotypes that endure throughout adulthood in chimpanzees. Interestingly, a recent cross taxa meta-analysis found a generally stronger influence of maternal effects on trait variation in general in species *without* parental care compared to those *with* parental care (Moore et al., 2019). This meta-analysis included a number of studies on non-human primates and other mammal species in which postnatal care is present. However, none of these species has the extended period of immature dependency on mothers that is observed in human and non-human apes. Therefore, we anticipated both genetic and non-genetic maternal effects to strongly contribute to variation in this phenotype in chimpanzees.

## Results

We used long-term behavioral, demographic, and physiological data collected between 2000 and 2018 from two field sites of two sub-species of chimpanzee. In Taï National Park (5°52’N, 7°20’E), Côte d’Ivoire, data were collected from three communities of western chimpanzees (East, North, and South; Wittig and Boesch, 2019) and in Budongo Conservation Field Station, Uganda (2°03′N,31°46’E), data were collected from two communities of eastern chimpanzees (Sonso and Waibira; Reynolds, 2005; Samuni et al., 2014).

Urine and fecal samples were collected from individuals of all ages (2-53 years old) within these communities. For each urine sample (n=6,123 samples), we quantified cortisol levels using liquid chromatography-tandem mass spectrometry (LCMS; Hauser et al., 2008) and corrected for variation in water content in the urine using the specific gravity (SG) of each sample (Miller et al., 2004). Therefore, we report urinary cortisol levels as ng cortisol/ml SG. From the fecal samples, we genotyped DNA extracts using a two-step amplification method including 19 microsatellite loci (per Arandjelovic et al., (2009).

In combination with behavioral observations of mother-offspring dyads, these genotypes allowed us to generate a pedigree containing 159 named mothers and 50 named fathers; 310 offspring had known mothers and 185 offspring had both known mothers and fathers). Following stringent criteria to measure circadian cortisol responses (see below), we included 170 individuals from this pedigree in our final dataset. Table 1 describes sampling by pedigree and group. Figure S1 in the Supplementary Materials illustrates the pedigree for individuals with urinary cortisol values in our study.

**Table 1:**
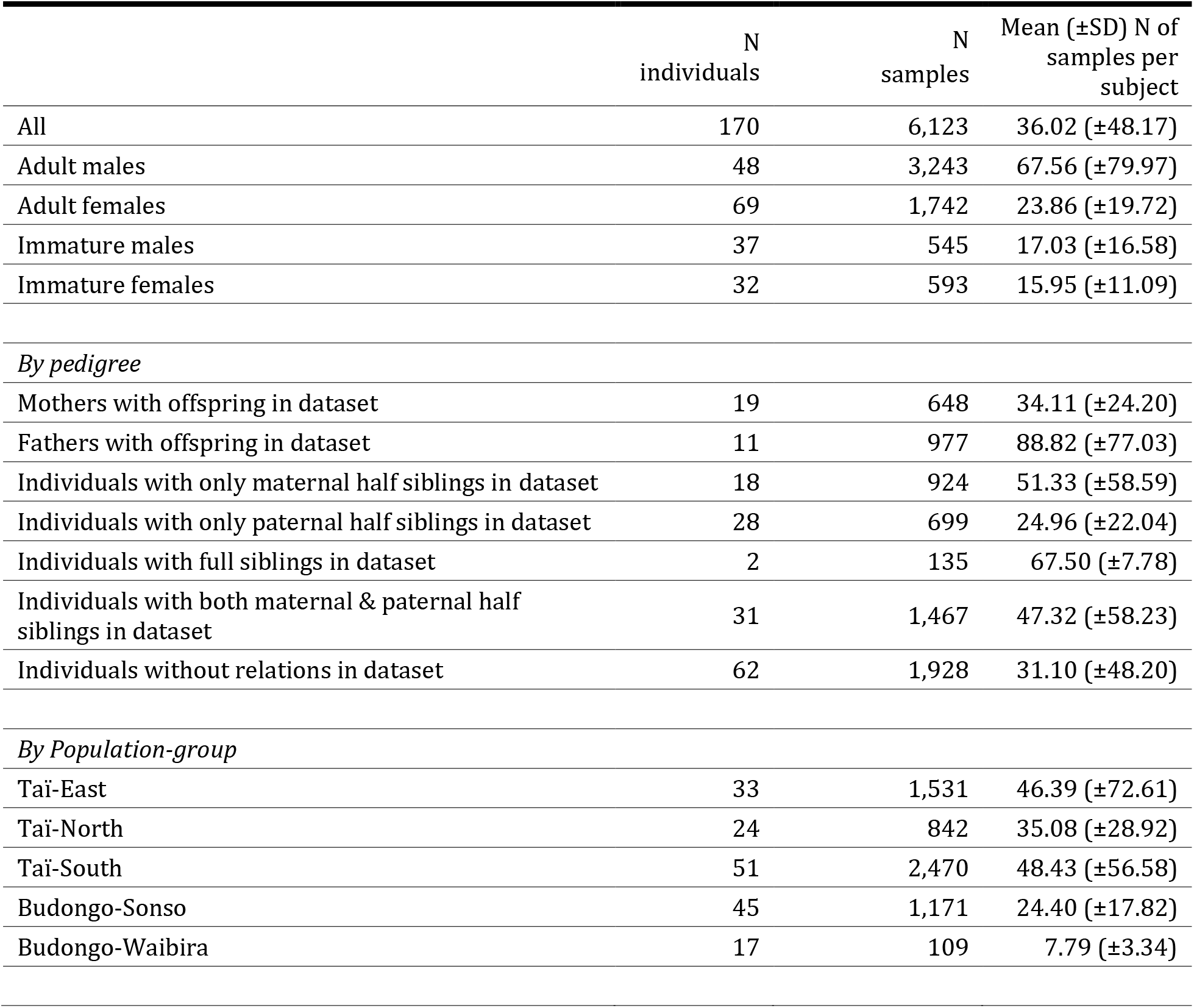
Summary statistics for final dataset used in the study. In total, 6,123 urinary cortisol values from 170 individuals were included in the study. Note that certain individuals fall into several pedigree categories (e.g. an individual can be a father and have a maternal or paternal sibling), therefore, the number of individuals in pedigree categorization exceeds 170. The range of numbers of years of sampling of individuals in the dataset was 1-13 years, with a mean ± SD of 2.63 ± 3.01 years.

| | N<br>individuals | N<br>samples | Mean (±SD) N of<br>samples per<br>subject |
| --- | --- | --- | --- |
| All | 170 | 6,123 | 36.02 (±48.17) |
| Adult males | 48 | 3,243 | 67.56 (±79.97) |
| Adult females | 69 | 1,742 | 23.86 (±19.72) |
| Immature males | 37 | 545 | 17.03 (±16.58) |
| Immature females | 32 | 593 | 15.95 (±11.09) |
| *By pedigree* |  |  |  |
| Mothers with offspring in dataset | 19 | 648 | 34.11 (±24.20) |
| Fathers with offspring in dataset | 11 | 977 | 88.82 (±77.03) |
| Individuals with only maternal half siblings in dataset | 18 | 924 | 51.33 (±58.59) |
| Individuals with only paternal half siblings in dataset | 28 | 699 | 24.96 (±22.04) |
| Individuals with full siblings in dataset | 2 | 135 | 67.50 (±7.78) |
| Individuals with both maternal & paternal half siblings in dataset | 31 | 1,467 | 47.32 (±58.23) |
| Individuals without relations in dataset | 62 | 1,928 | 31.10 (±48.20) |
| *By Population-group* |  |  |  |
| Tai-East | 33 | 1,531 | 46.39 (±72.61) |
| Tai-North | 24 | 842 | 35.08 (±28.92) |
| Tai-South | 51 | 2,470 | 48.43 (±56.58) |
| Budongo-Sonso | 45 | 1,171 | 24.40 (±17.82) |
| Budongo-Waibira | 17 | 109 | 7.79 (±3.34) |

### Repeatability

We used linear mixed-effect models (LMMs) with a Gaussian error structure to test adjusted repeatability, i.e., the proportion of variance attributable to between-individual differences given conditional effects (Dingemanse and Dochtermann, 2013; Nakagawa and Schielzeth, 2010), of both urinary cortisol levels (*R*^2^) and cortisol reaction norms (*RN*^2^), i.e. circadian cortisol responses. Our key predictor of cortisol level variation (log-transformed to achieve a symmetrical distribution) was time of day, which we converted into a continuous, hours-since-midnight value for each sample. Previous research found higher *RN*^2^ for the quadratic term of time of day in our study populations (Sonnweber et al., 2018); therefore, we included time of day as both linear and quadratic terms to model the potential circadian responses. We included as fixed effects variables previously shown to influence urinary cortisol levels (see Materials & Methods for full details): the age of the individual at the day of sampling (in years); group size; the male-to-female sex-ratio; the sine and cosine of date (to account for seasonality); LCMS methodology; and a categorical variable delineating individuals based on demography and reproductive state (five levels: “adult male”, “lactating female”, “cycling female”, “immature male”, and “immature female”; see Methods for description of variables). For the random effects of all models, we created a factor variable composed of group identity and the sampling year (termed “group-year”), and a variable to account for samples being pooled from various research projects (“project identity”).

We fitted three models: (i) an *intercept null* model, which included the fixed and random effects described above, (ii) a *random intercept* model, which added random intercepts for individual identity and a dummy variable composed of individual identity and the sampling year (termed “ID-year”; used to compare within-year and between year repeatability, see below), and (iii) a *reaction norm* model by including random slopes for the linear and quadratic terms of time of day within the random effects of individual identity and ID-year.

Using a model comparison approach and leave-one-out cross validation (Vehtari et al., 2021, 2019, 2017), we found strong support for the inclusion of the random intercepts for individual identity, but weak support for the inclusion of random slopes within these effects (Table S1). This pattern was also reflected in the observed repeatability estimates (Table 2). Using custom code adapted from a previous study (Sonnweber et al., 2018), from the *reaction norm* model, we calculated a within-year *R*^2^ estimate (variance explained by the ID-year variable) of 0.09 (95% confidence intervals = 0.06, 0.13) and a between-years *R*^2^ estimate (individual identity variable variance) of 0.05 (95% confidence intervals = 0.02, 0.07). We found substantial support for consistent individual differences in circadian reaction norm intercepts, i.e., average cortisol levels given the effect of time of day, with a *RN*^2^ estimate for the intercept of 0.47 (95% confidence intervals = 0.30, 0.67). Although the mean *RN*^2^ estimates for the linear and quadratic time of day slopes, 0.20 and 0.21 respectively, suggested a substantial proportion of variance in these phenotypes are attributed to individual differences, these estimates were associated with a large amount of uncertainty, with the lower credible intervals of both slopes close to 0.

**Table 2:** Repeatability coefficients from reaction norm models quantifying circadian cortisol responses in wild chimpanzees. Repeatability coefficients were calculated across all individuals (n=170), then within the specific demographics of adult males (n=46), adult females (n=69), and immatures (n=69). Note that certain individuals (n=14) appear both as adults and immatures in the overall dataset.

| Demographic | Coefficient | Estimate | (lCI, uCI) |
| --- | --- | --- | --- |
| All individuals combined | Within-year $R^2$ | 0.09 | 0.06, 0.13 |
| | Between years $R^2$ | 0.05 | 0.02, 0.07 |
| | $RN^2$ intercept | 0.47 | 0.30, 0.67 |
| | $RN^2$ linear slope | 0.20 | 0.00, 0.86 |
| | $RN^2$ quadratic slope | 0.21 | 0.00, 0.89 |
| Adult males | Within-year $R^2$ | 0.08 | 0.04, 0.14 |
| | Between years $R^2$ | 0.04 | 0.00, 0.08 |
| | $RN^2$ intercept | 0.44 | 0.13, 0.77 |
| | $RN^2$ linear slope | 0.27 | 0.00, 0.91 |
| | $RN^2$ quadratic slope | 0.25 | 0.00, 0.93 |
| Adult females | Within-year $R^2$ | 0.06 | 0.00, 0.13 |
| | Between years $R^2$ | 0.05 | 0.00, 0.11 |
| | $RN^2$ intercept | 0.87 | 0.61, 0.99 |
| | $RN^2$ linear slope | 0.47 | 0.00, 0.99 |
| | $RN^2$ quadratic slope | 0.39 | 0.00, 0.98 |
| Immatures | Within-year $R^2$ | 0.20 | 0.08, 0.33 |
| | Between years $R^2$ | 0.08 | 0.00, 0.18 |
| | $RN^2$ intercept | 0.43 | 0.00, 0.79 |
| | $RN^2$ linear slope | 0.22 | 0.00, 0.74 |
| | $RN^2$ quadratic slope | 0.19 | 0.00, 0.69 |

The apparent lack of between individual differences in circadian slopes was unexpected given the strong evidence for consistent individual differences in this phenotype in a previous study of adult male chimpanzees, a dataset which included individuals in our present study (Sonnweber et al., 2018). Therefore, to examine if the inclusion of adult females and immatures in our dataset contributed to uncertainty to our *RN*^2^ slope estimates, we ran repeatability analyses for each separate demographic (adult males, adult females, immatures; see Supplementary Materials for model specifications). For all demographics, we still observed a high amount of uncertainty for our *RN*^2^ slope estimates (Table 2). Generally, across and within demographics we found strong support for consistent individual differences in reaction norm intercepts rather than slopes. The *RN*^2^ intercept estimates for adult males and females were clearly non-zero (Table 2); for immatures, although the estimate was high (*RN*^2^ = 0.43), the CI range was very wide, suggesting uncertainty.

Figure 1 illustrates the urinary cortisol circadian responses of four randomly selected father-mother-offspring triads from four groups in our study (for the Waibira group, we had insufficient numbers of individuals to represent such a triad). Figures S3-S5 in the Supplementary Materials respectively illustrate the circadian cortisol responses for all adult male, adult female, and immature subjects included in the study. Tables S2-S9 in the Supplementary Materials provides the model summary for the fixed and random effects of the *reaction norm* models of each demographic.

**Figure 1:**
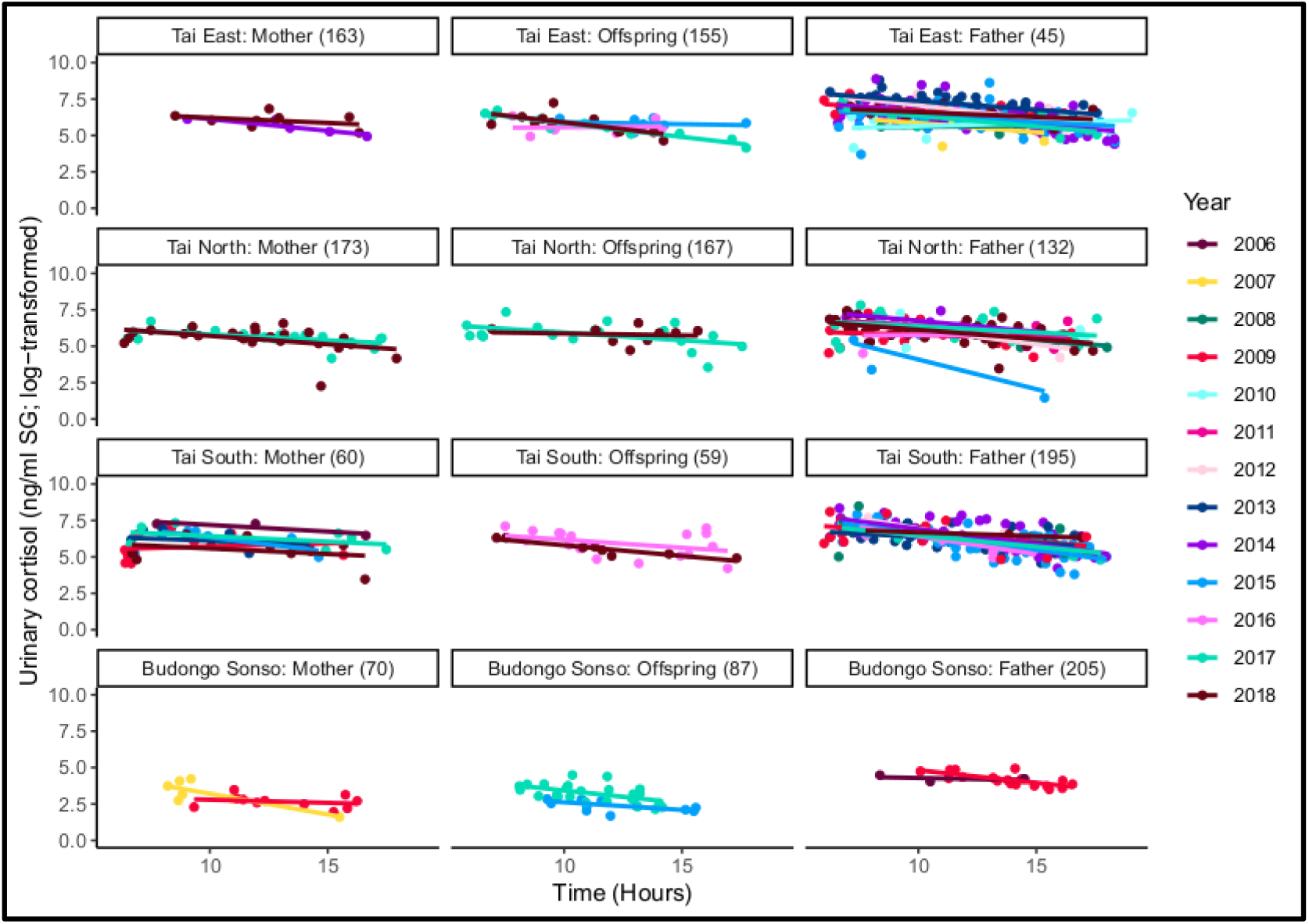
Circadian reaction norms for urinary cortisol levels (ng/ml SG; log transformed) for triads of father-mother-offspring in four of our study communities (for the Budongo Waibira group, we had insufficient numbers of individuals to represent such a triad). The points represent individual sample values, the slopes individual responses to time of day; both sample values and responses are shaded according to the year in which they were collected, respectively. The numbers in parentheses above each panel indicate the identity of the individual as it appears in the pedigree (Figure S1).

### Heritability

We estimated the heritability of urinary cortisol levels and circadian cortisol responses by implementing an “animal model” (Wilson et al., 2010), which estimates additive genetic variance in a trait. Our animal model was identical in structure to those constructed for repeatability, with the major exception being the inclusion of the pedigree as a random effect (Wilson et al., 2010). In addition, to partition the relative contribution of maternal effects (the main caregiver), we also included the identity of the mother of the individual sampled as a random effect.

We computed the genetic (h^2^) and maternal (m^2^) components of heritability as the proportion of inter-individual variance explained by the pedigree and the maternal identity, respectively. Specifically, we calculated h^2^ and m^2^ for the inter-individual variance in the average cortisol levels (h^2^_intercept_ and m^2^_intercept_), and in cortisol responses to the linear (h^2^_linear_ and m^2^_linear_) and quadratic (h^2^_quadratic_ and m^2^_quadratic_) terms for time of day. We also estimated the proportion of covariance between intercept, linear slope, and quadratic slopes explained by additive genetic or maternal factors (Wilson et al., 2010).

The relative contribution of our random effects to variation in circadian cortisol responses in wild chimpanzees are shown in Figure 2, with a summary of the maternal and genetic effects in Table 3 (full details of all variance components are in Table S10 and Table S11 of the Supplementary Materials). Maternal effects explain about 10% of the variance of the reaction norm intercept (m^2^_intercept_ = 0.10), with 90% credibility intervals (hereafter 90% CI, 0.02-018) higher than the point estimate for genetic effects, which is an order of magnitude lower (h^2^_intercept_ = 0.01). Specifically, we estimate that 93% of the probability mass of m^2^_intercept_ is higher than the posterior probability of m^2^_intercept_. For the linear and quadratic circadian slope terms, the 90% CIs of the proportion of variance explained by maternal effects are very wide (m^2^_linear_ = 0.09; 0.000-0.51; m2quadratic = 0.03; 0.00-0.27) and prevent a comparison with that explained by genetic effects (h^2^_linear_ = 0.06; 0.00-0.42; h^2^_quadratic_ = 0.04; 0.00-0.36). Similar estimates were obtained using independent models, in which either group identity was used as predictor in place of continuous predictors such as group size (Figure S6, Table S12), or in which only individuals sampled in the Taï forest (4,843 samples belonging to 111 individuals) were used, excluding the possibility that artifacts due to unaccounted population structure are present (Figure S7, TableS13).

**Table 3:** Summary of genetic (h^2^) and maternal (m^2^) effect estimates on circadian cortisol responses in wild chimpanzees. Each coefficient represents a different component of the circadian cortisol response. We also report the proportion of permutations for which these coefficient estimates were larger than in the observed data. Coefficients in bold were larger in our observed data than in at least 95% of our random permutations.

| Coefficient | Estimate | (lCI, uCI) | Proportion<br>observed <<br>permutations |
| --- | --- | --- | --- |
| *Genetic effect* |  |  |  |
| $h^2_{\text{intercept}}$ | 0.01 | (0.00, 0.06) | 0.92 |
| $h^2_{\text{linear}}$ | 0.06 | (0.00, 0.42) | 0.84 |
| $h^2_{\text{quadratic}}$ | 0.04 | (0.00, 0.36) | 0.90 |
| *Maternal effect* |  |  |  |
| **$m^2_{\text{intercept}}$** | **0.10** | **(0.02, 0.18)** | **0.00** |
| $m^2_{\text{linear}}$ | 0.09 | (0.00, 0.51) | 0.22 |
| $m^2_{\text{quadratic}}$ | 0.03 | (0.00, 0.27) | 0.06 |

**Figure 2:**
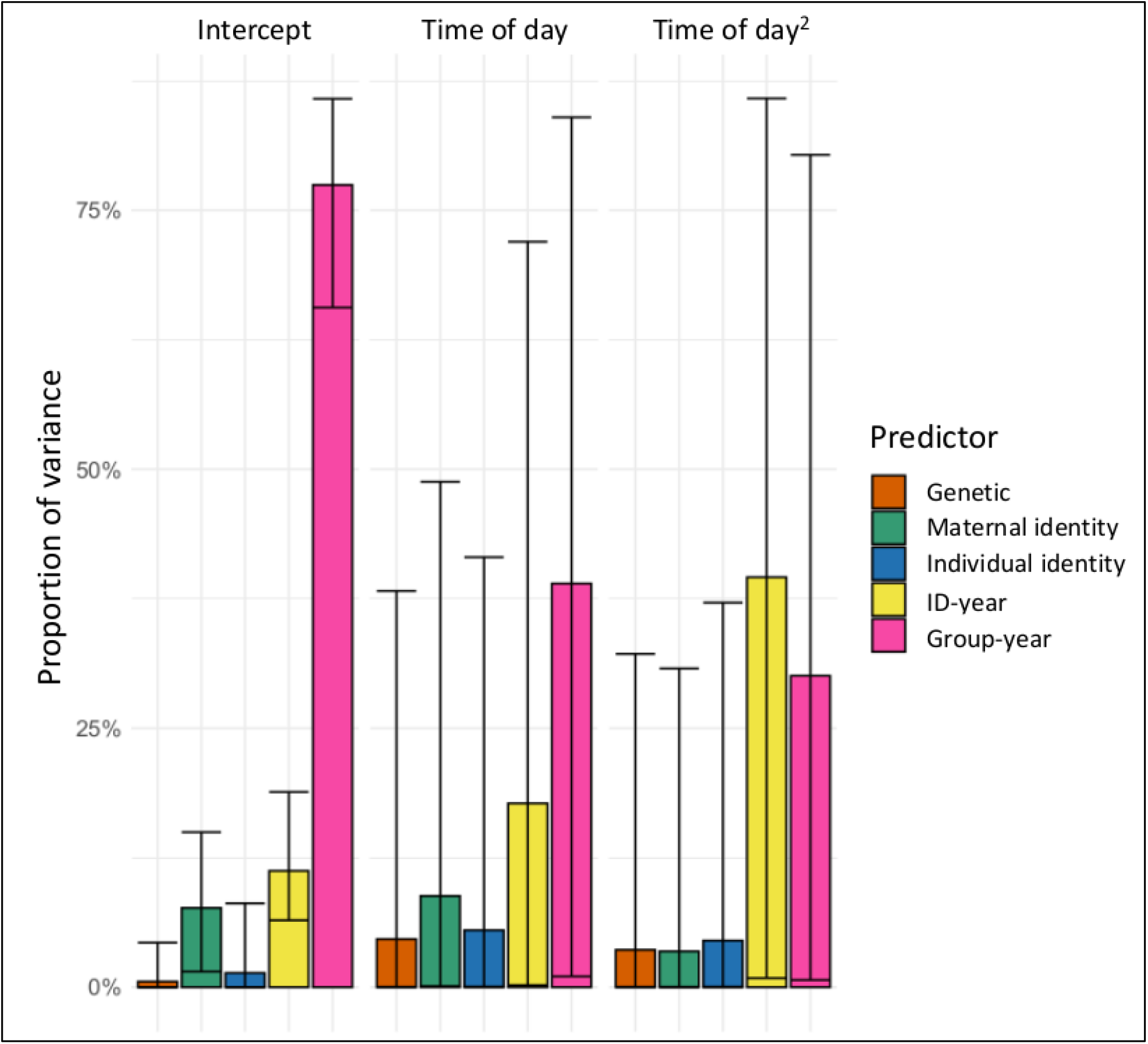
Estimates for the proportion of variance among the random effects in our model examining variation in circadian cortisol responses in wild chimpanzees. The error bars represent the 95% credible interval range of the estimates.

Note, the CIs of our h^2^ and m^2^ estimates indicate a large degree of uncertainty (see Table 3). In addition, their values are by definition bound to be positive as they are derived from the variance components of the random effects in the animal model. Hence, to assess whether maternal and genetic factors determine detectable non-zero effects and to test whether the differences between m^2^ and h^2^ could be due to chance, we performed re-sampling of the data and calculated the proportion of cases in which estimates were higher than for the observed data (i.e., false positives). Specifically, we reshuffled the identities of the individuals within their communities (and thus maintaining control of group-level environmental and social factors) 100 times in the additive genetic matrix. Individuals newly classified as siblings after the permutation of the genetic matrix, were assigned to the same mother in the predictor “maternal identity”, so that genetic relationships and maternal effects were always concordant. By doing this, we obtained permutations of the data that simulated genetic and maternal relationships expected by chance, while leaving unaltered the effects of all other predictors, keeping the same structure in the additive genetic matrix, and the same distribution of maternal relationships among individuals (Figure 3).

**Figure 3:**
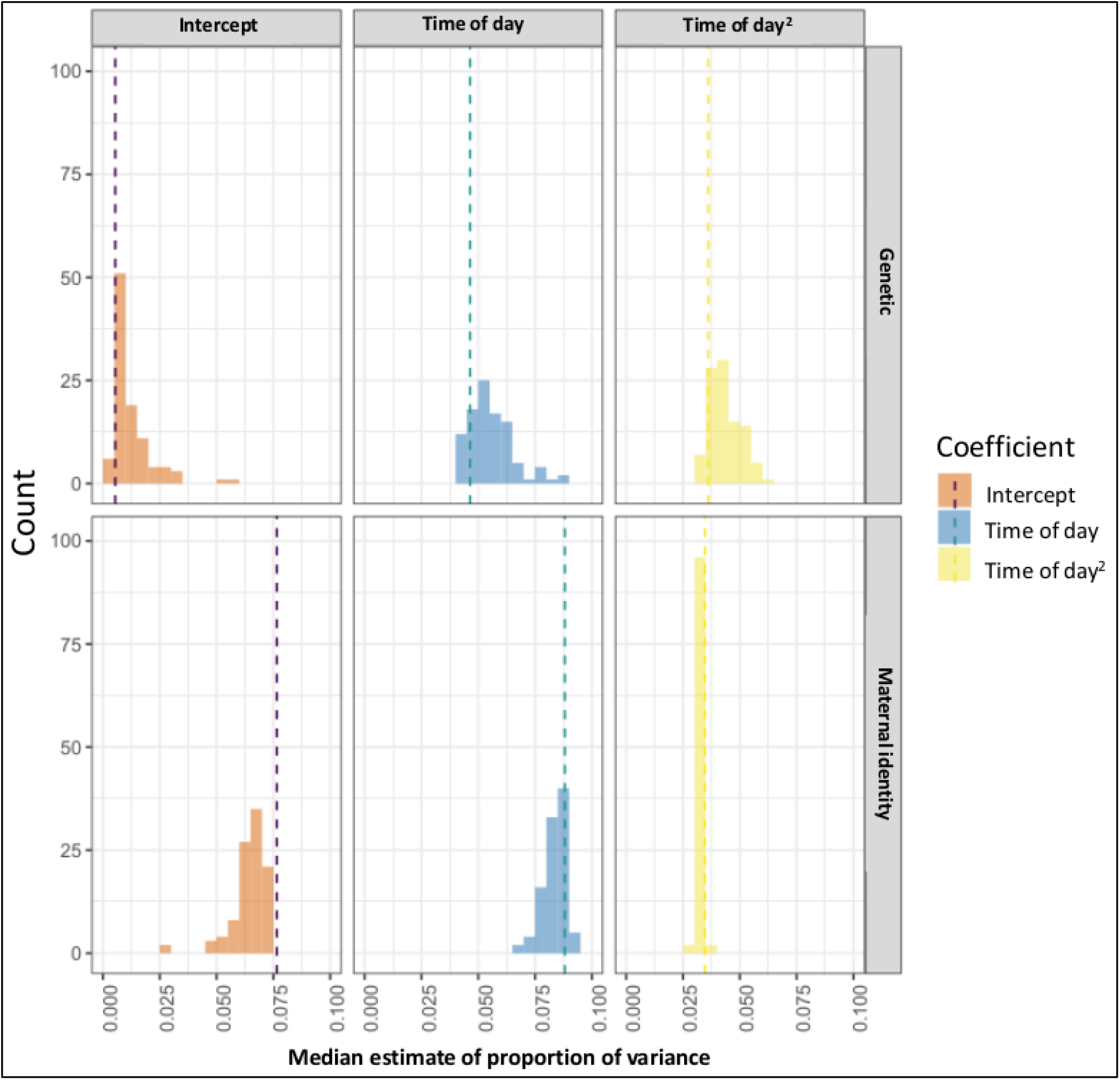
Median proportion of variance estimates obtained from the observed data (dashed vertical lines) versus estimates obtained by 100 datasets with permuted genetic relationships between individuals. Histograms represent the counts of each estimate value from the permutations. In this permutation analysis, the proportion of variance calculations includes all random effects, including our technical predictor, “project identity”. Our final reported maternal effect estimate is higher than presented here as we consider only the biological predictors in that calculation. Figure S9 in the supplementary materials illustrates the permutations of all variance components in our heritability model.

For m^2^_intercept_, all permutations had estimates lower than the observed data (Table 3; Figure 3), suggesting that the observed effects cannot be explained by chance. The same pattern was replicated when group was included in the model instead of group size or only a single site was used (Table S12, S13). These results confirm a non-zero contribution of maternal effects to the cortisol phenotypes of wild chimpanzees. All other observed coefficients of genetic or non-genetic maternal effects were in the same range as those derived from random permutations (Table 3).

We also used permutations to test whether the observed difference between the variance explained by the maternal and genetic effects can occur because of chance alone. None of the permutations indicated a higher difference between the variance explained by maternal and genetic effects than those observed in the data in either the model with group size (Figure 4) or the model with community included as a predictor (Figure S8). We conclude that the maternal environment is more influential than genetics in shaping cortisol responses in wild chimpanzees.

**Figure 4:**
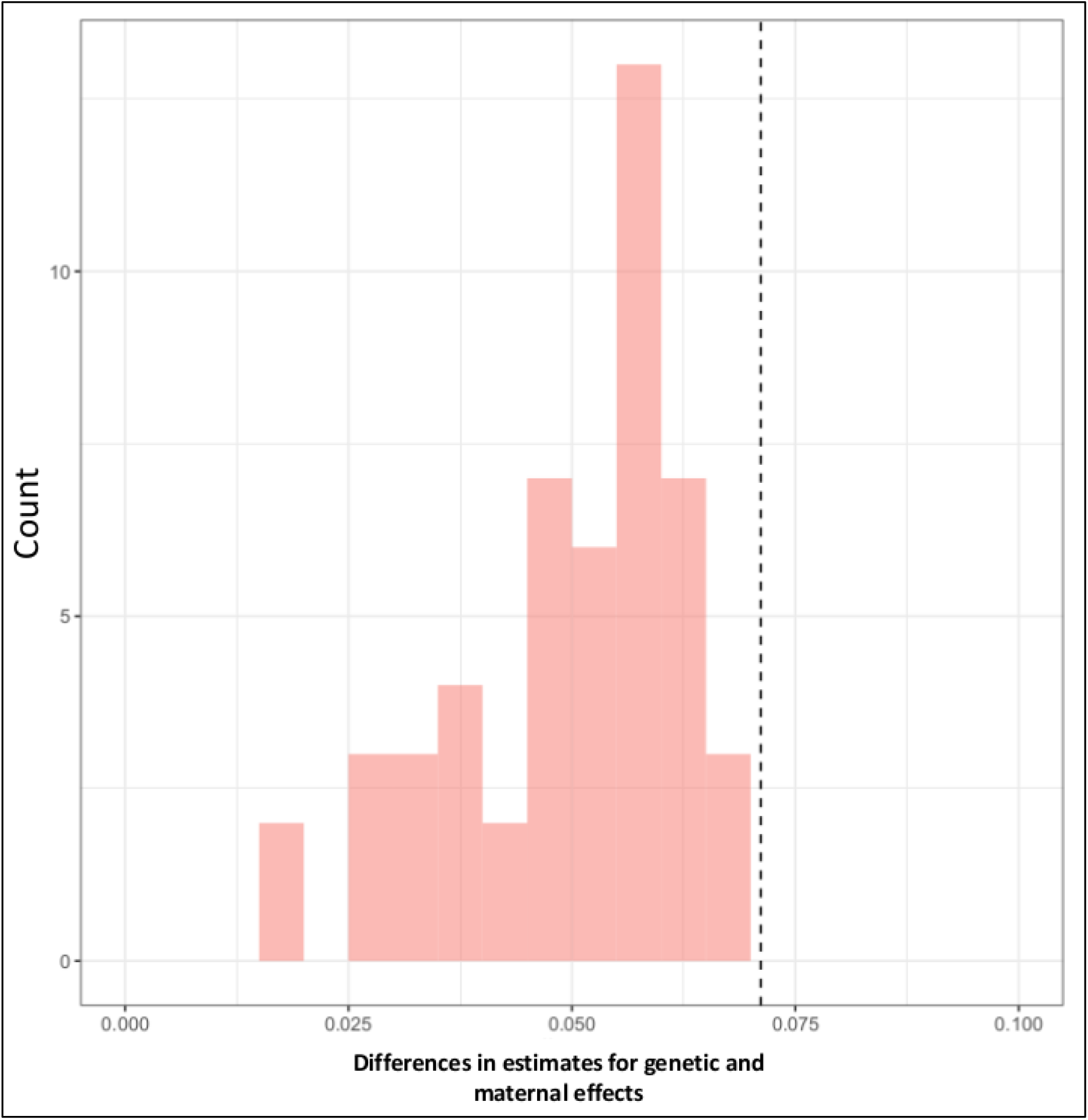
Estimates of the difference in the proportion of variance explained by the maternal effect and that explained by genetic factors in the observed data (dashed line) and in 100 permutations of the data (red histogram).

## Discussion

Our study leverages almost two decades of long-term data collection of more than 6,000 urine samples from 170 individuals to identify consistent individual differences in circadian cortisol responses in wild chimpanzees. Using this unique dataset, we find that the maternal environment has a primary role in shaping circadian cortisol phenotypes in this species, certainly when compared to the influence of genetic factors. Our results are robust to different model structures and are corroborated by permutations of the data which indicate our maternal and genetic effect estimates are not artefacts of group structures. Our study shows the importance of long-term data collection in the wild, especially for long-lived species, and raises important biological questions about the nature of the non-genetic maternal effects we documented. We estimated that ~10% of variation in average cortisol levels (conditional on the effect of time of day), is due to these maternal factors.

In our study, much of the variation in urinary cortisol levels was attributable to short-term group-level (“group-year” random effect) and individual-level (“ID-year” random effect) factors. This finding illustrates the flexibility of these phenotypes in wild animals, which vary due to food availability (Wessling et al., 2018a), dominance hierarchy instability (Preis et al., 2019), reproductive state (Emery Thompson et al., 2010) or age (Emery Thompson et al., 2020). While we attempted to control for such factors (see Materials and Methods), our non-invasive and non-experimental approach inherently contributed unidentifiable confounds that might well explain variation in cortisol phenotypes. For example, short-term elevations in cortisol levels can occur in chimpanzees following single aggressive encounters (Wittig et al., 2015) or disturbance from neighboring communities of conspecifics (Samuni et al., 2019), and inter-individual differences exist in the magnitude of these elevations depending on the amount of social support available to them (Wittig et al., 2016). Given these potential confounds, it is notable that we were able to identify such a clear maternal effect in our study. Although absence of evidence is not evidence of absence, the lack of a clear genetic effect in our results at least indicates a qualitatively stronger influence of maternal identity in shaping cortisol phenotypes in our study population. In a recent meta-analysis, Moore et al (2019) found a limited role for parental care in shaping the strength of parental effects on trait variation. However, as far as we are aware, few species included in the study demonstrate the prolonged mother-offspring association observed in chimpanzees. We hope that our results will encourage studies in other animals with protracted developmental phases or maternal associations to compare and contrast the relative influence of mothers and genetic inheritance.

Determining the specific mechanism leading to the observed effect of the maternal environment, such as protracted maternal care or epigenetic processes, merits further study. Chimpanzees have slow life histories, characterized by long gestation and maturation relative to lifespan (Bründl et al., 2021), as well as prolonged dependency on maternal care (Crockford et al., 2020; Nakamura et al., 2014; Samuni et al., 2020; Stanton et al., 2020). Recent evidence suggests that adult female chimpanzees, and thus mothers, have both relatively stable dominance hierarchies compared to males (Mielke et al., 2019) and consistent individual differences in social phenotypes that endure over several years (Tkaczynski et al., 2020b). As offspring associate almost permanently with their mothers until around the age of 12 years (Reddy and Sandel, 2020), maternal social phenotype is the key determinant of the social environment of immature offspring. As some mothers are more consistently gregarious than others (Tkaczynski et al., 2020b), and social settings likely impact rates of exposure to social stressors for offspring (Sabbi et al., 2021; Tkaczynski et al., 2020a), maternal effects on average cortisol levels are perhaps not surprising in immature individuals. However, the maternal effects observed in our study apply for individuals of all age classes, suggesting that they endure beyond the immature phase.

Based on their dominance rank and social phenotypes, mothers likely vary in their ability to secure feeding resources for their offspring, and this may have lifelong consequences for the growth and the foraging skills of their offspring (Estienne et al., 2019; Samuni et al., 2020). Previous research suggests maternal dominance status influences fecal GC levels in male, but not female immature chimpanzees (Murray et al., 2018), therefore, status alone is unlikely to explain the full extent of the maternal effect identified in our study. If mothers vary in their rates of direct social interaction with their offspring and others, via grooming or food sharing for example, offspring may learn variable social or technical skills, such as extractive foraging (Estienne et al., 2019). The stable social phenotypes observed in adult chimpanzees includes rates of aggression, with some individuals being consistently more aggressive than others over the lifespan (Tkaczynski et al., 2020b). Therefore, long-term mother-offspring association may also behaviorally prime offspring on how to deal with social antagonism or other social challenges. As chimpanzees are long-lived, offspring that remain in their natal group (i.e., all males and a small percentage of females) may even inherit certain social relationships or components of their mother’s social networks (Langergraber et al., 2013). Therefore, maternal effects may influence the social and ecological environment of offspring throughout their life, as well as prime how they react to these environments on a physiological and behavioral level.

In rodents, early life adversity, such as maternal neglect or loss, can induce hyper-methylation of DNA regions coding for GC receptors, leading to lifelong alterations in the sensitivity of these receptors and thus affecting GC feedback loops and overall GC levels (Champagne and Curley, 2009; Zhang et al., 2013). In long-lived primates, including in humans, early life adversity can lead to long-term alteration of HPA axis activity (Berens et al., 2017; Ehrlich et al., 2016; Rosenbaum et al., 2020). Indeed, in wild baboons, early life adversity can have intergenerational effects on survival, i.e., if a mother experiences adversity, both she and her offspring can experience reduced survival outcomes (Zipple et al., 2019), which may be explained by the GC effects of adversity. Recent meta-analyses and evidence from long-term field studies now suggest that, at least for long-lived species, elevated HPA axis activity over the lifespan is a predictor of survival (Bonier et al., 2009; Campos et al., 2021; Schoenle et al., 2021). However, in wild chimpanzees, although maternal loss impacts later life reproductive success (Crockford et al., 2020), there is no evidence that this is the result of long-term HPA axis activity alteration as effects on circadian cortisol patterns following maternal loss do not endure into adulthood (Girard-Buttoz et al., 2021). This time-limited nature of alteration of the HPA axis activity suggests that adversity may not have a clear epigenetic effect on HPA axis activity in this species, at least among young orphan individuals that later survive into adulthood. Whether the enduring maternal effect observed in our study is due to early life epigenetic maternal effects, or whether it is the result of the aforementioned behavioral priming, will not be trivial to disentangle. Behavioral observations can help determine whether mother-offspring dyads and maternal siblings are exposed to similar levels of social stressors, or whether the same dyads and siblings behaviorally respond to stressors in a similar way. Although this would not eliminate the possibility of epigenetic effects, it would allow empirical testing for evidence of behavioral priming.

In our study, the contribution of heritable factors to cortisol phenotypes was low as compared to values reported in human twin studies (e.g. 60%; Gustafsson et al., 2011), and more controlled laboratory (e.g. 28%; Houslay et al., 2019) or wild experimental animal studies (e.g. 40%; Bairos-Novak et al., 2018) in which GC variation was directly manipulated by the observers. Indeed, our analysis revealed an extremely low and unstable estimate of the contribution of genetics to variation in chimpanzee cortisol phenotypes, contrary to our predictions. Human research involves more controlled sampling than can be achieved with wild animals, especially when non-invasive and non-experimental methods are used, as in our study. To address this methodological challenge, we employed strict criteria for the inclusion of individuals into the study to ensure we could accurately characterize their cortisol phenotypes. We required that each individual have at least one year of sampling in which we had samples spanning the majority of the day (i.e., morning, midday, and evening samples) in order to measure circadian responses and their repeatability. Employing such criteria reduced the number of individuals we could include in the study, and all individuals were spread across five separate groups and two different populations (note that we repeated our analysis solely within the larger of these two populations, finding qualitatively similar results despite the reduced overall sample size; Table S12 in Supplementary Materials). Chimpanzees are also a long-lived species with low fertility. Consequently, despite working with data from two of the longest running wild chimpanzee field sites (Reynolds, 2005; Wittig and Boesch, 2019), our pedigree is relatively shallow for this form of analysis, including relatively few third-generation individuals. Despite these challenges, our study reveals new insights on how cortisol phenotypes vary across different demographics of wild chimpanzees, and the prominent role of maternal effects in shaping these differences.

Previous studies examining the repeatability of circadian cortisol responses in chimpanzees focused exclusively on adult males (Sonnweber et al., 2018); in our study we were able to show that individual circadian cortisol responses are repeatable across demographics, including adult females in various reproductive states and in immature individuals. However, we only found strong support only for consistent individual differences in average cortisol levels, rather than circadian slopes. This difference from the findings in Sonnweber et al. (2018) was not driven by the inclusion of adult females and immature individuals, as in our separate adult male repeatability analysis, we again found weak support for consistent individual differences in circadian slopes. Within our adult male only analysis, as compared to Sonnweber et al. (2018), we included substantially more samples and individuals, despite using stricter criteria for individual inclusion. Circadian slopes vary with experiences of adversity, including maternal loss and illness (Behringer et al., 2020; Girard-Buttoz et al., 2021), and also change with aging and life history stages in chimpanzees (Emery Thompson et al., 2020). Therefore, our uncertain repeatability estimates for circadian slopes could be due to substantial within-individual variation. Given our study included only healthy chimpanzees and modelled age effects, it seems more likely that our uncertain repeatability estimates for slopes are the result of low between-individual variation for this particular component of circadian cortisol phenotypes.

To conclude, in our study, the maternal environment is the main early life influence on cortisol regulation throughout the lifespan in chimpanzees. Whether this is due to epigenetic processes early in development, or due to behavioral priming of how to deal with the ecological or social environment, clearly merits further investigation and will contribute to our understanding of the role of parents and developmental plasticity in long-lived species. Indeed, determining whether this maternal effect on cortisol regulation has been specifically selected for, or is instead a by-product of extended maternal association, will be key to understanding prolonged development and parental dependency as a life history adaptation.

## Materials & Methods

### Study Site & Subjects

In both Taï and Budongo, data on the chimpanzees are systematically collected by a combination of locally-employed field assistants and visiting researchers. Longitudinal data includes daily counts of group compositions, as well as recording of behavioral and social interactions using a combination of focal observations and ad-libitum sampling (Altmann, 1974). During observations of the chimpanzees, observers opportunistically collected urine and fecal samples from identifiable individuals. In Taï, regular observations of the chimpanzees commenced in 1990 (North, 1990-present; South, 1999-present; East, 2007-present (Wittig and Boesch, 2019)) and regular urine sample collection (see below) commenced in 2000 (North and South, 2000-present; East, 2003-present). In Budongo, regular observations of the chimpanzees commenced in 1994 (Sonso, 1994-present; Waibira, 2011-present; (Reynolds, 2005; Samuni et al., 2014)) and regular urine sample collection commenced in 2005 (Sonso, 2005-present; Waibira, 2017-present).

### Urine Sample Collection and Analysis

We collected urine from identifiable individuals using a plastic pipette to transfer urine from the ground or vegetation into a 5 ml cryovial. Cryovials were stored in liquid nitrogen once back in camp, typically within 12 hours of collection. Frozen samples were transported packed in dry ice to the Max Planck Institute for Evolutionary Anthropology in Leipzig, Germany, where they were stored at ≤20°c in freezers.

We quantified urinary cortisol levels for each sample using LCMS ((Hauser et al., 2008)) and MassLynx (version 4.1; QuanLynx-Software). We used prednisolone (coded as “old method” in models, i.e. most samples analyzed prior to July 2016; Hauser et al., 2008), or testosterone d4 (“new method”, i.e. all samples analyzed post September 2016; Wessling et al., 2018b) as the internal standards. For each sample, we measured specific gravity (SG) using a refractometer (TEC, Ober-Ramstadt, Germany). SG values were used to correct cortisol measurements for variation in water content in the urine using the formula outlined by Miller et al. (2004):

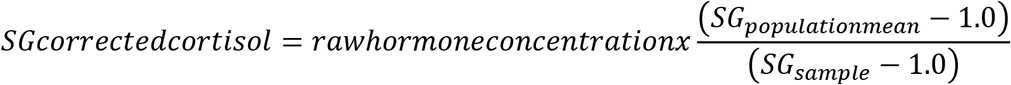

The population means were derived from the samples included in this analysis. The SG population mean was 1.02 for Taï and 1.02 for Budongo.

### Fecal Sample Collection and Pedigree Generation

Fecal samples were collected from identifiable individuals. The samples were collected using plastic bags and then either directly stored in ethanol, dried on silica gel, or using a two-step ethanol-silica method (Nsubuga et al., 2004). Dried samples were transported in silica to the Max Planck Institute for Evolutionary Anthropology in Leipzig, Germany. Approximately 100mg of each sample was extracted using either the QIAamp DNA stool (Qiagen) or the GeneMATRIX Stool DNA Purification (Roboklon) kits. We genotyped DNA extracts using a two-step amplification method including 19 microsatellite loci as detailed previously (Arandjelovic et al., 2009). Using CERVUS 3.0 software (Kalinowski et al., 2007), we compared the resultant genotypes using the ‘identity analysis’ function to confirm individual identities and the ‘parentage analysis’ function to confirm maternities and assign paternities.

### Data Preparation

To provide an accurate measure of circadian patterns for each individual, we excluded certain samples where cortisol levels were expected to be elevated and not representative of normal circadian patterning. Here, we provide a detailed description of the sample exclusion process.

In female primates, including chimpanzees, cortisol levels vary with reproductive state (Brent et al., 2011; Cohen et al., 1958; Emery Thompson et al., 2010). Chimpanzee gestation is approximately 240 days (Peacock and Rogers, 1959). Using demography data and the birth dates of offspring, we assigned females to three reproductive states (Emery Thompson et al., 2010): pregnant (during the 240 days preceding the birth of any offspring), lactating (the 1,095 days [based on average resumption of cycling in the population] subsequent to the birth of any offspring) and cycling (any other period of time when females were not assigned as pregnant or lactating). We included all adult female samples where we were able to assign reproductive state to the female at the time of sampling (Kahlenberg et al., 2008). Furthermore, following related studies (Emery Thompson et al., 2020, 2010), we excluded samples from pregnant females because cortisol levels tend to increase during pregnancy. In fact, interactions can occur between maternal and fetal HPA axes making it difficult to accurately determine maternal cortisol levels in isolation (Smith and Thomson, 1991).

In immature chimpanzees (<12 years old), maternal separation elevates cortisol secretion and has short-term effects on cortisol circadian patterns (Girard-Buttoz et al., 2021). Therefore, if immature individuals lost their mother prior to the age of 12 years old (social maturity), we excluded any sample collected from them following maternal loss during immaturity. However, as there is no evidence of long-term impacts of maternal loss in mature chimpanzees (Girard-Buttoz et al., 2021), all mature individuals were included regardless of maternal loss during immaturity. Furthermore, injury and sickness can elevate cortisol levels in primates (Barton, 1987; Behringer et al., 2020; McIntosh, 1987; Muehlenbein and Watts, 2010) and affect circadian cortisol patterns in chimpanzees (Behringer et al., 2020). Therefore, we excluded samples from individuals that displayed symptoms of sickness or injury (determined by onsite veterinarians in each field site).

Lastly, there is a link between dominance rank and GC levels in male and female chimpanzees (Markham et al., 2014; Muller and Wrangham, 2004). However, for one group in our study (Waibira), we had insufficient data to calculate ranks for the females, and in all groups, for immature individuals it is unclear whether maternal rank influences their cortisol levels. Given these caveats, we did not assign ranks or include this as a variable in our analyses when combing demographics. However, when we analyzed repeatability in the demographics separately, for the adult male analysis, we included dominance rank as a fixed effect in those models. Male dominance ranks were calculated using pant grunt vocalizations, a unidirectional call given from subordinate individuals (Wittig and Boesch, 2003). We used a likelihood-based adaptation of the Elo rating approach to calculate ranks (Foerster et al., 2016; Mielke et al., 2018; Neumann et al., 2011); we assigned continuous Elo ranks to subjects for each day of sampling; each score was standardized between 0 (lowest rank) and 1 (highest rank) within each group. By pooling males, females, and immature individuals together without the inclusion of dominance rank in our heritability analysis, our estimates of heritable contributions to those differences are likely more conservative.

To ensure that we were able to characterize circadian cortisol patterns for each individual, we only included individuals with a minimum of 3 urine samples per year, collected during both morning and afternoon hours, such that the earliest and latest samples were separated by at least 6 hours.

To accurately model circadian patterns of cortisol for all individuals (our measure of cortisol reaction norm), we included interactions between the linear and quadratic time variables and all other fixed effects. We used 12 years of age to distinguish between adult (aged >=12 years) and immature individuals (aged <12 years), as it is the age at which individuals socialize and forage predominantly independent from their mothers (Reddy and Sandel, 2020). In addition to the demographic categorization (adult male, cycling female, lactating female, immature male, immature female) and age of each individual on the day of sampling, we included in the analysis a number of control variables known to influence cortisol levels. Both group size and mating competition (Emery Thompson et al., 2010; Muller and Wrangham, 2004; Preis et al., 2019; Samuni et al., 2019) can affect GC levels in primates, therefore, we calculated both the number of adults (mean[+SD]; East 13.81[+2.19], North 8.92[+1.33], South 16.52[+2.58], Sonso 36.35[+4.12], Waibira 54.11[+2.50]) and the male-to-female sex-ratio (mean[+SD]; East 0.35[+0.12], North 0.50[+0.19], South 0.37[+0.10], Sonso 0.49[+0.05], Waibira 1.02[+0.02]) at the time of sampling for each sample. Lastly, as seasonal variation in rainfall, temperature, humidity and food availability can influence cortisol levels in chimpanzees (Wessling et al., 2018a), we accounted for this circannual variation by converting the Julian date of sampling into a circular variable and including its sine and cosine in our models (Stolwijk et al., 1999; Wessling et al., 2018a, 2018b).

### Notes on Model Fitting and Verification

All data preparation, models and analyses were performed using R version 3.6.1 (R Core Team, 2020). Prior to testing our models, we applied the *vif* function of the ‘car’ R package (Fox and Weisberg, 2011) to linear model versions of our mixed models (i.e. lacking random effects) to test for any collinearity issues via examination of variance inflation factors (VIF). There were issues with collinearity if either “site” or “group” were included in the models as both variables were either collinear with each other or with “group size”. Therefore, we retained just “group size”, with all remaining VIFs < 2.90. The “group-year” variable was also included as a random effect to account for group-level confounds. Furthermore, for the heritability analyses, we performed additional analyses using models containing “group” as a predictor, finding no qualitative differences in our animal model estimates (see Table S12).

All models were fitted with a Gaussian error distribution using the R package ‘brms’ (Hadfield, 2019). For all models, numeric variables were standardized as z-scores. We fit models with weakly regularising priors for the fixed effects (β~Normal(0,1)) and for the random effects (student t-distributed (3, 0, 10)), with uniform (LKJ(1)) priors for covariance matrices of the random slopes. For all models, we specified four chains of 4,000 iterations, half of which were devoted to the warm-up. Sampling diagnostics (Rhat < 1.1) and trace plots confirmed chain convergence for all models. Effective sample sizes confirmed no issues with autocorrelation of sampling for all models.

We estimated the heritability of urinary cortisol levels and their circadian patterning by fitting an “animal model”, which estimates additive genetic variance in a trait by including the pedigree of individuals as a random effect (Wilson et al., 2010). Pedigrees were generated with the R package ‘MasterBayes’ (Hadfield, 2017). The additive genetic matrix was computed using the Amatrix function of the R package ‘AGHmatrix’ (Amadeu et al., 2016).

## Data availability

All data used in the analyses presented are available via Figshare (https://doi.org/10.6084/m9.figshare.13720765.v1).

## Acknowledgments

We thank the Ministère de l’Enseignement Supérieur et de la Recherche Scientifique, the Ministère de Eaux et Fôrests in Côte d’Ivoire, the Office Ivoirien des Parcs et Réserves, the Uganda Wildlife Authority and the Uganda National Council for Science and Technology for permitting the study. In Côte d’Ivoire, we are grateful to the Centre Suisse de Recherches Scientifiques en Côte d’Ivoire and the staff members of the Taï Chimpanzee Project for their support. In Uganda, we thank the management and staff of the Budongo Conservation Field Station. We are indebted to the efforts of Christophe Boesch and Vernon Reynolds in the establishments of the study field sites and their contributions to years of data collection. We also thank the many field and research assistants that help generate the data for this project. We are extremely grateful for the work conducted in the laboratories of Tobias Deschner and Linda Vigilant in the Max Planck Institute of Evolutionary Anthropology, Leipzig, Germany, specifically the efforts of Róisín Murtagh, Vera Schmeling, Janette Gleiche, Anette Nicklisch, Juliane Damm, Carolyn Rowney, and Jared Cobain. We also thank Ruth Sonnweber and Verena Behringer for useful discussions on the topic.

## Author Contributions

CC, CGB, FM and PJT conceived the study. AP, CC, CG, CGB, CYA, EGW, LS, LW, PF, PJT, PDV, RMW, TD, TL, VM, and ZS collected data. TD, LS, CH, KZ, CC, and RMW provided long-term data. PJT, FM, CC, CGB, PF, TD and RMW helped design the study; FM, CGB and PT performed the statistical analyses; TD oversaw the laboratory analyses; LV supervised and conducted genetic parentage analyses; PT wrote the first draft of the manuscript, all authors contributed to subsequent editing.

