## Supplementary material for "Non-genetic maternal effects shape individual differences in cortisol phenotypes in wild chimpanzees": Supplemtary Tables & Figures

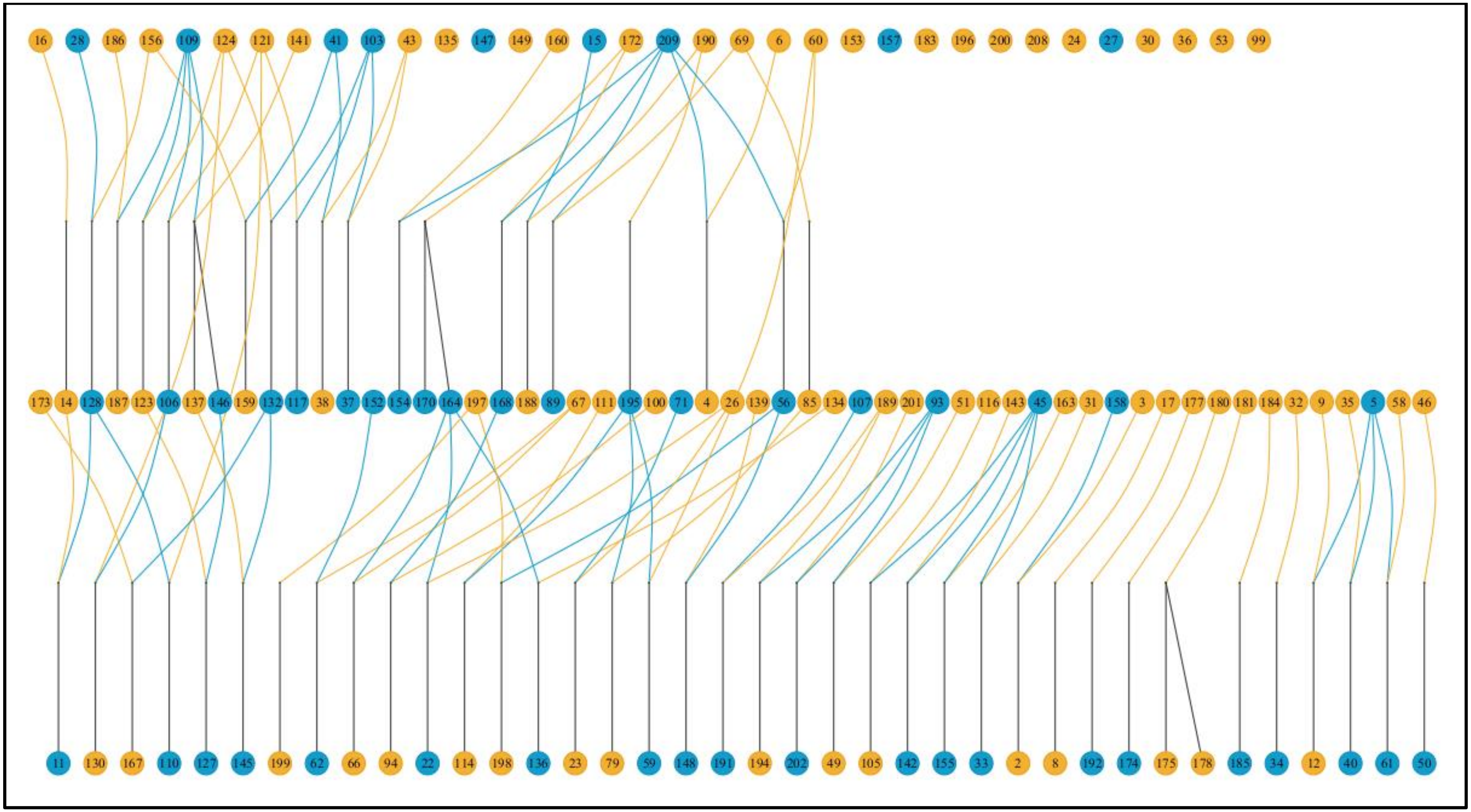


*Figure S1: Pedigree of individuals from the Taï field site included within our analysis. Males are represented by the blue circles, females by the yellow circles. Note, this pedigree illustration includes the mothers and/or fathers of individuals with urinary cortisol values in our study, even if these mothers and/fathers were not sampled themselves. However, the offspring of individuals are only included if the offspring themselves had urinary cortisol values in the study.*

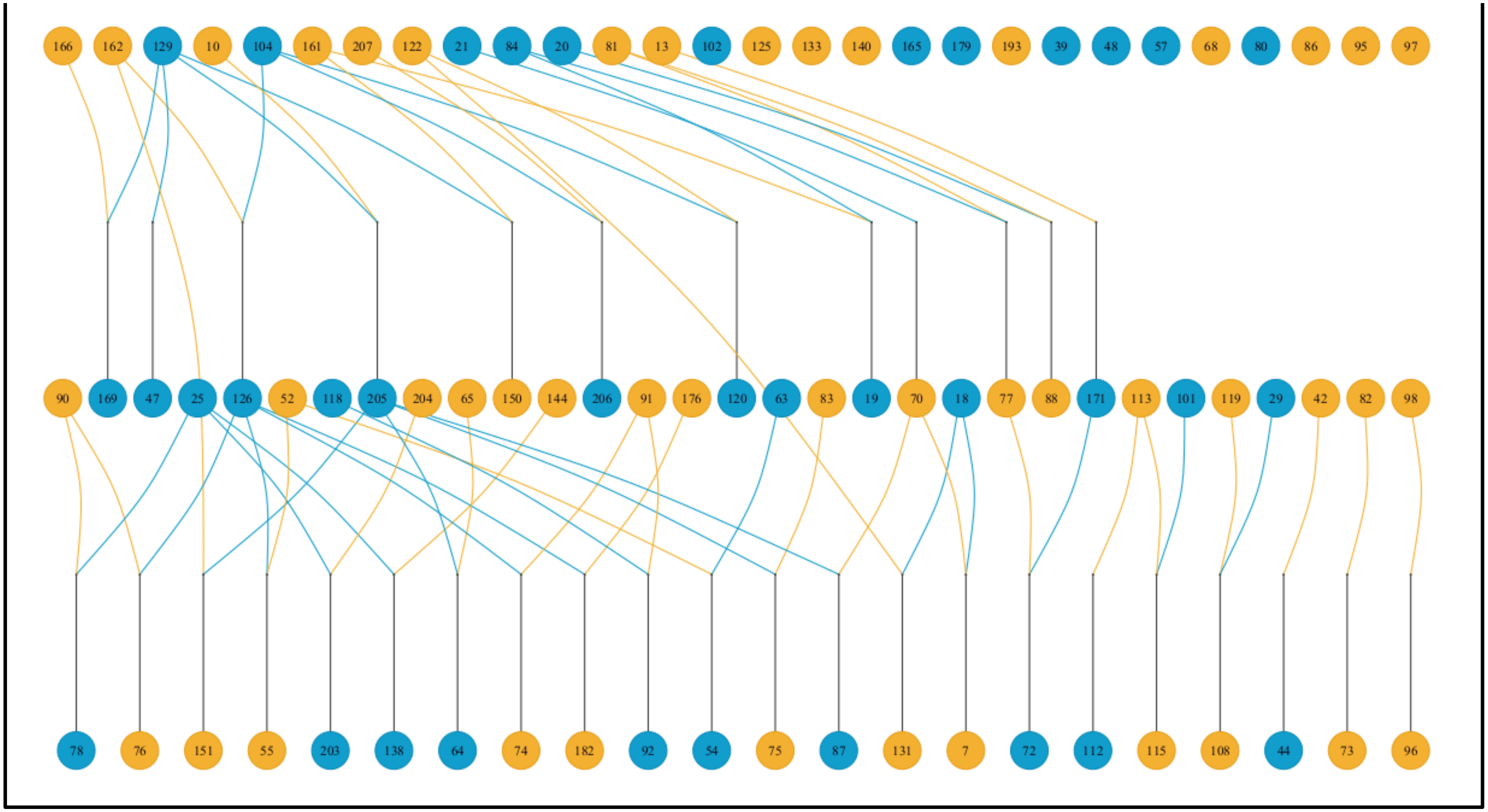


*Figure S2: Pedigree of individuals from the Budongo field site included within our analysis. Males are represented by the blue circles, females by the yellow circles. Note, this pedigree illustration includes the mothers and/or fathers of individuals with urinary cortisol values in our study, even if these mothers and/fathers were not sampled themselves. However, the offspring of individuals are only included if the offspring themselves had urinary cortisol values in the study.*

*Table S1: Model comparison results using leave-one-out cross validation and the loo_compare function of the 'loo' R package. Model comparison were conducted for all individuals, then for models built for adult males, adult females, and juvenile individuals separately. In each case, the model with the strongest support is highlighted in bold.*

| Demographic | Model | Expected logwise predictive density | Standard error |
|---|---|---|---|
| All individuals | **Random intercept** | **0.000** | **0.000** |
| | Reaction norm | -2.076 | 4.635 |
| | Null model | -300.966 | 25.831 |
| Adult males | **Random intercept** | **0.000** | **0.000** |
| | Reaction norm | -2.767 | 3.649 |
| | Null model | -95.016 | 13.999 |
| Adult females | **Random intercept** | 0.000 | 0.000 |
| | Reaction norm | -1.070 | 2.444 |
| | Null model | -60.092 | 12.048 |
| Juveniles | **Reaction norm** | **0.000** | **0.000** |
| | Random intercept | -13.715 | 7.823 |
| | Null model | -62.516 | 13.767 |

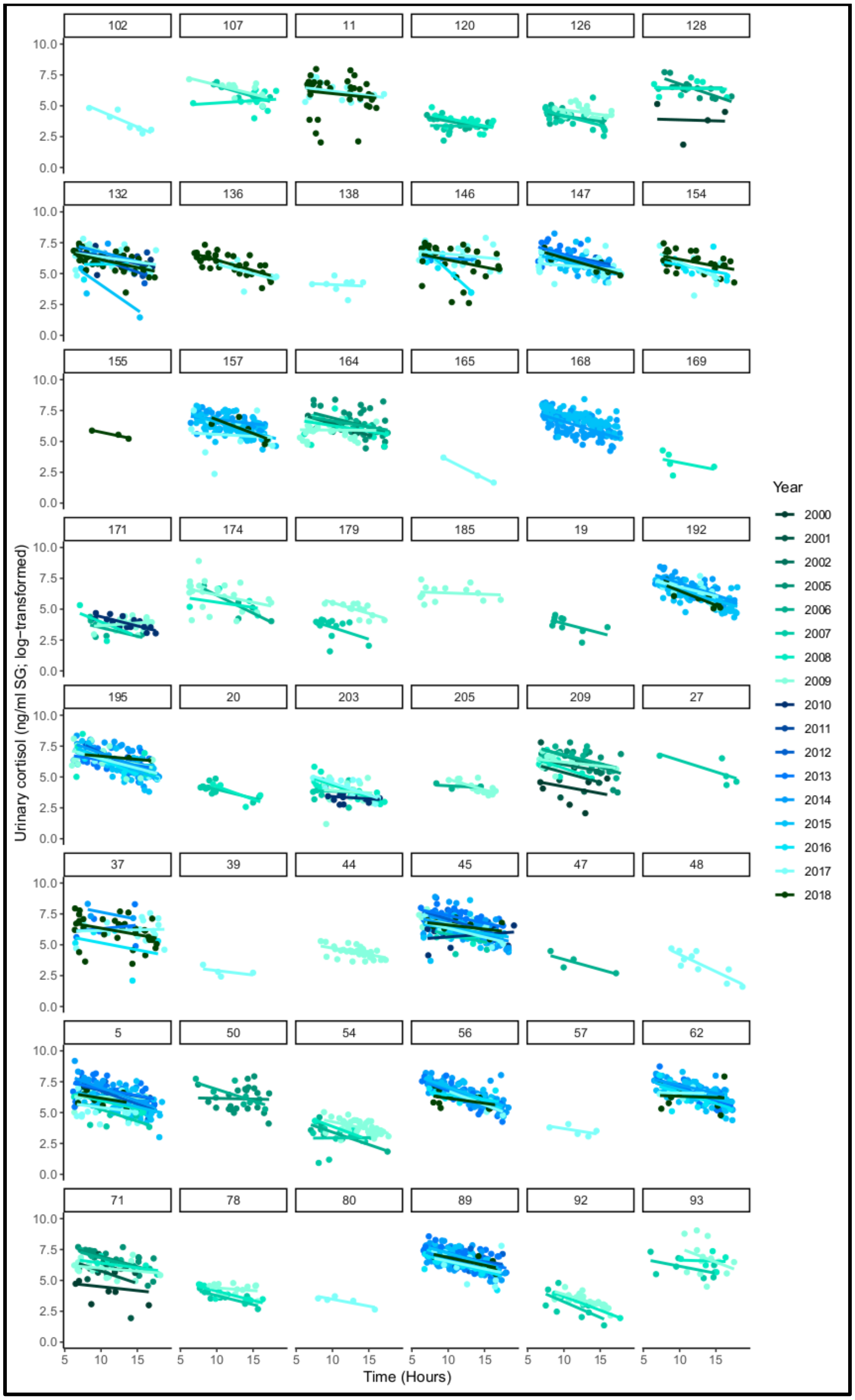


*Figure S3: Urinary cortisol concentration (ng/ml SG; log transformed) circadian reaction norms for all individuals when appearing as adult males in the study (n =48). The points represent individual sample values, the slopes individual responses to time of day; both sample values and responses are shaded according to the year in which they were collected respectively. The numbers above panels indicate individual identity as it appears in the pedigree (Figure S1).*

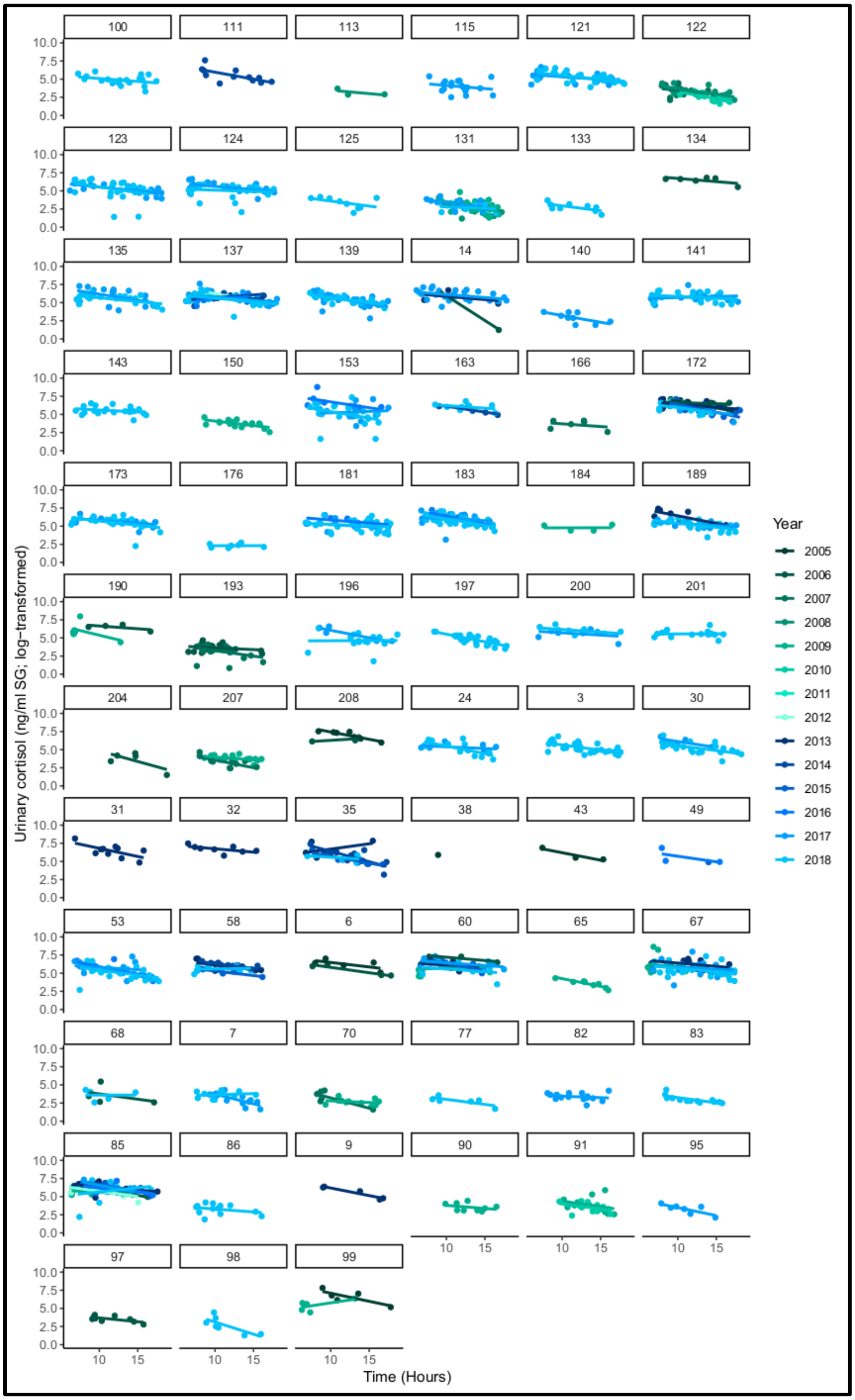


*Figure S4: Urinary cortisol concentration (ng/ml SG; log transformed) circadian reaction norms for all individuals when appearing as adult females in the study (n =69). The points represent individual sample values, the slopes individual responses to time of day; both sample values and responses are shaded according to the year in which they were collected respectively. The numbers above panels indicate individual identity as it appears in the pedigree (Figure S1).*

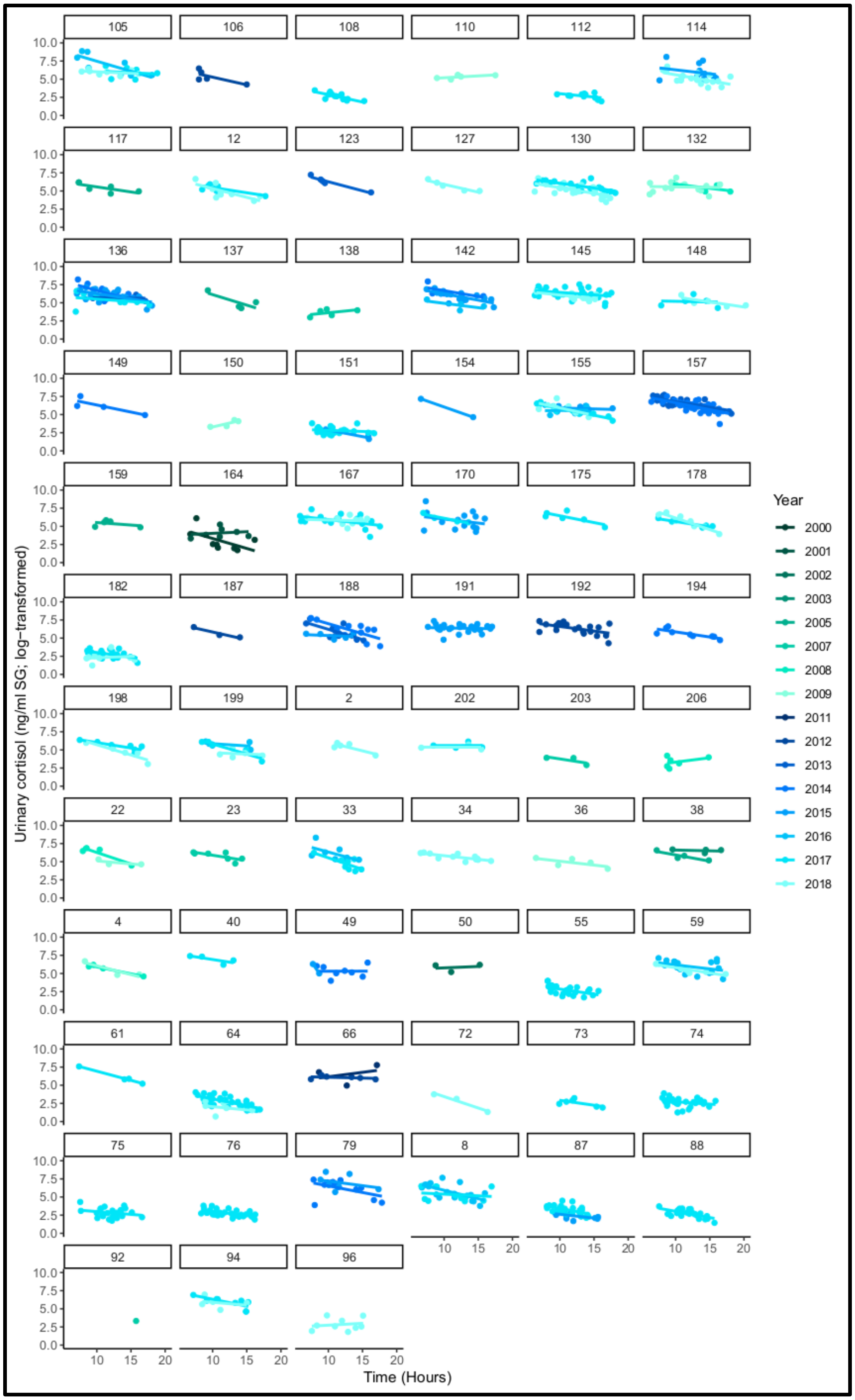


*Figure S5: Urinary cortisol concentration (ng/ml SG; log transformed) circadian reaction norms for all individuals when appearing as immature individuals in the study (n = 69). The points represent individual sample values, the slopes individual responses to time of day; both sample values and responses are shaded according to the year in which they were collected respectively. The numbers above panels indicate individual identity as it appears in the pedigree (Figure S1).*

*Table S2: Fixed effect results from the reaction norm LMM estimating urinary cortisol concentrations (ng/ml SG) in response to time of day in five chimpanzee communities. Categorical variables have the reference category in parentheses. Effects where 95% credible intervals do not cross 0 are indicated in bold.*

| Variable | Estimate | Est. error | Q2.5 | Q97.5 |
|---|---|---|---|---|
| Intercept | 4.911 | 0.195 | 4.501 | 5.280 |
| Time of day$^2$ | -0.182 | 0.218 | -0.602 | 0.254 |
| **Time of day** | **-0.617** | **0.119** | **-0.861** | **-0.392** |
| Demographic (adult males) | | | | |
| **Cycling female** | **0.422** | **0.114** | **0.201** | **0.653** |
| Juvenile female | -0.106 | 0.119 | -0.339 | 0.130 |
| Juvenile male | -0.097 | 0.128 | -0.346 | 0.151 |
| Lactating female | -0.023 | 0.127 | -0.270 | 0.228 |
| **Age at sample** | **0.227** | **0.068** | **0.091** | **0.361** |
| **Sin date** | **-0.133** | **0.021** | **-0.174** | **-0.092** |
| Cos date | 0.121 | 0.022 | 0.078 | 0.164 |
| Sex ratio | 0.078 | 0.160 | -0.238 | 0.399 |
| **Community size** | **-1.410** | **0.302** | **-1.944** | **-0.782** |
| **LCMS method (new method)** | **0.396** | **0.159** | **0.089** | **0.719** |
| Time of day$^2$: Demographic (adult males) | 0.067 | 0.223 | -0.380 | 0.493 |
| Time of day$^2$: Cycling female | -0.202 | 0.238 | -0.670 | 0.242 |
| Time of day$^2$: Juvenile female | 0.198 | 0.255 | -0.296 | 0.700 |
| Time of day$^2$: Lactating female | 0.188 | 0.254 | -0.316 | 0.668 |
| Time of day$^2$: Age at sample | -0.109 | 0.107 | -0.328 | 0.093 |
| Time of day$^2$: Sin date | 0.042 | 0.058 | -0.073 | 0.160 |
| **Time of day$^2$: Cos date** | **0.162** | **0.058** | **0.048** | **0.273** |
| Time of day$^2$: Sex ratio | -0.156 | 0.101 | -0.354 | 0.041 |
| Time of day$^2$: Community size | 0.152 | 0.116 | -0.080 | 0.373 |
| Time of day$^2$: LCMS method (new method) | -0.155 | 0.107 | -0.373 | 0.053 |
| Time of day: Demographic (adult males) | -0.092 | 0.121 | -0.319 | 0.152 |
| Time of day: Cycling female | -0.100 | 0.128 | -0.340 | 0.162 |
| Time of day: Juvenile female | -0.132 | 0.137 | -0.398 | 0.131 |
| Time of day: Lactating female | -0.026 | 0.135 | -0.285 | 0.247 |
| **Time of day: Age at sample** | **0.151** | **0.055** | **0.043** | **0.259** |
| **Time of day: Sin date** | **-0.088** | **0.027** | **-0.142** | **-0.035** |
| Time of day: Cos date | -0.126 | 0.029 | -0.181 | -0.069 |
| Time of day: Sex ratio | 0.050 | 0.046 | -0.041 | 0.140 |
| Time of day: Community size | -0.058 | 0.053 | -0.161 | 0.045 |
| Time of day: LCMS method (new method) | -0.084 | 0.053 | -0.187 | 0.024 |

*Table S3: Random effect results from the reaction norm LMM estimating urinary cortisol concentrations (ng/ml SG) in response to time of day in five chimpanzee communities.*

| Groups | Name | Variance | Est.error | Q2.5 | Q97.5 |
|---|---|---|---|---|---|
| ID-year | Intercept | 0.23 | 0.03 | 0.18 | 0.28 |
| | Time of day$^2$ | 0..19 | 0.10 | 0.01 | 0.38 |
| | Time of day | 0.13 | 0.05 | 0.01 | 0.22 |
| Individual identity | Intercept | 0.21 | 0.03 | 0.16 | 0.27 |
| | Time of day$^2$ | 0.07 | 0.05 | 0.00 | 0.20 |
| | Time of day | 0.04 | 0.03 | 0.00 | 0.11 |
| Group-year | Intercept | 0.61 | 0.09 | 0.45 | 0.80 |
| Database code | Intercept | 0.43 | 0.17 | 0.20 | 0.85 |
| Residual | | 0.65 | 0.01 | 0.64 | 0.66 |

*Table S4: Fixed effect results from the reaction norm LMM estimating urinary cortisol concentrations (ng/ml SG) in response to time of day among adult males in five chimpanzee communities. Categorical variables have the reference category in parentheses. Effects where 95% credible intervals do not cross 0 are indicated in bold.*

| **Variable** | **Estimate** | **Est. error** | **Q2.5** | **Q97.5** |
|---|---|---|---|---|
| Intercept | 5.651 | 0.17 | 5.298 | 5.968 |
| Time of day$^2$ | -0.194 | 0.131 | -0.449 | 0.062 |
| **Time of day** | **-0.686** | **0.064** | **-0.810** | **-0.559** |
| **Rank** | **0.293** | **0.075** | **0.147** | **0.441** |
| **Age at sample** | **0.157** | **0.083** | **0.001** | **0.328** |
| **Sin date** | **-0.150** | **0.031** | **-0.208** | **-0.089** |
| **Cos date** | **0.095** | **0.032** | **0.029** | **0.157** |
| Sex ratio | -0.061 | 0.154 | -0.364 | 0.239 |
| **Community size** | **-1.013** | **0.241** | **-1.460** | **-0.516** |
| LCMS method (new method) | 0.133 | 0.184 | -0.214 | 0.517 |
| Time of day: Rank | 0.021 | 0.080 | -0.136 | 0.184 |
| Time of day: Age at sample | 0.128 | 0.065 | -0.003 | 0.252 |
| **Time of day: Sin date** | **-0.151** | **0.039** | **-0.227** | **-0.076** |
| **Time of day: Cos date** | **-0.173** | **0.041** | **-0.256** | **-0.091** |
| Time of day: Sex ratio | 0.050 | 0.064 | -0.074 | 0.174 |
| Time of day: Community size | -0.065 | 0.092 | -0.237 | 0.118 |
| Time of day: LCMS method (new method) | -0.111 | 0.075 | -0.259 | 0.039 |
| Time of day$^2$: Rank | 0.056 | 0.165 | -0.271 | 0.382 |
| Time of day$^2$: Age at sample | -0.129 | 0.127 | -0.372 | 0.124 |
| Time of day$^2$: Sin date | 0.038 | 0.085 | -0.126 | 0.202 |
| **Time of day$^2$: Cos date** | **0.271** | **0.089** | **0.095** | **0.452** |
| Time of day$^2$: Sex ratio | -0.182 | 0.134 | -0.452 | 0.087 |
| Time of day$^2$: Community size | 0.246 | 0.190 | -0.119 | 0.620 |
| Time of day$^2$: LCMS method (new method) | -0.070 | 0.151 | -0.365 | 0.221 |

*Table S5: Random effect results from the reaction norm LMM estimating urinary cortisol concentrations (ng/ml SG) in response to time of day among adult males in five chimpanzee communities.*

| **Groups** | **Name** | **Variance** | **Est.error** | **Q2.5** | **Q97.5** |
|---|---|---|---|---|---|
| ID-year | Intercept | 0.21 | 0.04 | 0.14 | 0.29 |
| | Time of day | 0.12 | 0.06 | 0.01 | 0.24 |
| | Time of day$^2$ | 0.20 | 0.12 | 0.01 | 0.44 |
| Individual identity | Intercept | 0.18 | 0.05 | 0.09 | 0.30 |
| | Time of day | 0.06 | 0.04 | 0.00 | 0.16 |
| | Time of day$^2$ | 0.08 | 0.06 | 0.00 | 0.23 |
| Group-year | Intercept | 0.62 | 0.10 | 0.45 | 0.83 |
| Database code | Intercept | 0.28 | 0.13 | 0.12 | 0.61 |
| Residual | | 0.66 | 0.01 | 0.64 | 0.68 |

*Table S6: Fixed effect results from the reaction norm LMM estimating urinary cortisol concentrations (ng/ml SG) in response to time of day among adult females in five chimpanzee communities. Categorical variables have the reference category in parentheses. Effects where 95% credible intervals do not cross 0 are indicated in bold.*

| **Variable** | **Estimate** | **Est. error** | **Q2.5** | **Q97.5** |
|---|---|---|---|---|
| Intercept | 5.001 | 0.356 | 4.272 | 5.717 |
| Time of day$^2$ | -0.133 | 0.275 | -0.678 | 0.404 |
| **Time of day** | **-0.543** | **0.133** | **-0.806** | **-0.286** |
| Reproductive state (cycling) | -0.095 | 0.127 | -0.343 | 0.157 |
| Age at sample | 0.159 | 0.098 | -0.035 | 0.351 |
| **Sin date** | **-0.148** | **0.041** | **-0.229** | **-0.068** |
| **Cos date** | **0.173** | **0.041** | **0.091** | **0.253** |
| Sex ratio | 0.008 | 0.137 | -0.289 | 0.261 |
| Community size | -0.669 | 0.594 | -1.875 | 0.381 |
| **LCMS method (new method)** | **0.620** | **0.306** | **0.051** | **1.240** |
| Time of day$^2$: Reproductive state (cycling) | -0.233 | 0.282 | -0.771 | 0.325 |
| Time of day$^2$: Age at sample | -0.024 | 0.175 | -0.374 | 0.313 |
| Time of day$^2$: Sin date | 0.156 | 0.112 | -0.061 | 0.385 |
| Time of day$^2$: Cos date | 0.034 | 0.111 | -0.179 | 0.256 |
| Time of day$^2$: Sex ratio | 0.033 | 0.229 | -0.420 | 0.489 |
| Time of day$^2$: Community size | 0.287 | 0.250 | -0.205 | 0.775 |
| Time of day$^2$: LCMS method (new method) | -0.418 | 0.226 | -0.866 | 0.016 |
| Time of day: Reproductive state (cycling) | -0.169 | 0.135 | -0.437 | 0.092 |
| **Time of day: Age at sample** | **0.171** | **0.081** | **0.012** | **0.329** |
| Time of day: Sin date | -0.029 | 0.053 | -0.132 | 0.074 |
| Time of day: Cos date | -0.069 | 0.052 | -0.174 | 0.031 |
| Time of day: Sex ratio | 0.137 | 0.098 | -0.056 | 0.331 |
| Time of day: Community size | -0.196 | 0.109 | -0.409 | 0.023 |
| Time of day: LCMS method (new method) | -0.061 | 0.107 | -0.270 | 0.149 |

*Table S7: Random effect results from the reaction norm LMM estimating urinary cortisol concentrations (ng/ml SG) in response to time of day among adult females in five chimpanzee communities.*

| **Groups** | **Name** | **Variance** | **Est.error** | **Q2.5** | **Q97.5** |
|---|---|---|---|---|---|
| ID-year | Intercept | 0.09 | 0.05 | 0.00 | 0.19 |
| | Time of day$^2$ | 0.23 | 0.13 | 0.01 | 0.48 |
| | Time of day | 0.08 | 0.06 | 0.00 | 0.21 |
| Individual identity | Intercept | 0.26 | 0.04 | 0.18 | 0.36 |
| | Time of day$^2$ | 0.16 | 0.11 | 0.01 | 0.40 |
| | Time of day | 0.07 | 0.05 | 0.00 | 0.17 |
| Group-year | Intercept | 0.17 | 0.07 | 0.04 | 0.33 |
| Database code | Intercept | 1.03 | 0.43 | 0.32 | 2.01 |
| Residual | | 0.66 | 0.01 | 0.63 | 0.68 |

*Table S8: Fixed effect results from the reaction norm LMM estimating urinary cortisol concentrations (ng/ml SG) in response to time of day among immatures in five chimpanzee communities. Categorical variables have the reference category in parentheses. Effects where 95% credible intervals do not cross 0 are indicated in bold.*

| **Variable** | **Estimate** | **Est. error** | **Q2.5** | **Q97.5** |
|---|---|---|---|---|
| Intercept | 4.801 | 0.204 | 4.404 | 5.198 |
| Time of day$^2$ | 0.102 | 0.158 | -0.201 | 0.412 |
| **Time of day** | **-0.836** | **0.077** | **-0.984** | **-0.683** |
| Sex (female) | -0.031 | 0.126 | -0.281 | 0.217 |
| Age at sample | -0.318 | 0.124 | -0.567 | -0.073 |
| Sin date | -0.061 | 0.047 | -0.157 | 0.030 |
| Cos date | 0.050 | 0.047 | -0.042 | 0.144 |
| Sex ratio | 0.080 | 0.137 | -0.191 | 0.351 |
| **Community size** | **-2.076** | **0.347** | **-2.728** | **-1.378** |
| LCMS method (new method) | 0.063 | 0.303 | -0.520 | 0.675 |
| Time of day$^2$: Sex (female) | 0.024 | 0.226 | -0.421 | 0.467 |
| Time of day$^2$: Age at sample | 0.236 | 0.225 | -0.197 | 0.698 |
| Time of day$^2$: Sin date | 0.016 | 0.137 | -0.251 | 0.289 |
| Time of day$^2$: Cos date | 0.235 | 0.129 | -0.024 | 0.487 |
| Time of day$^2$: Sex ratio | -0.211 | 0.211 | -0.632 | 0.204 |
| Time of day$^2$: Community size | -0.161 | 0.271 | -0.708 | 0.351 |
| Time of day$^2$: LCMS method (new method) | -0.408 | 0.374 | -1.141 | 0.315 |
| Time of day: Sex (female) | 0.022 | 0.116 | -0.212 | 0.245 |
| Time of day: Age at sample | -0.025 | 0.113 | -0.242 | 0.204 |
| Time of day: Sin date | 0.080 | 0.068 | -0.055 | 0.217 |
| Time of day: Cos date | -0.038 | 0.066 | -0.171 | 0.091 |
| Time of day: Sex ratio | 0.120 | 0.103 | -0.082 | 0.320 |
| Time of day: Community size | 0.164 | 0.126 | -0.084 | 0.410 |
| **Time of day: LCMS method (new method)** | **0.352** | **0.184** | **0.013** | **0.724** |

*Table S9: Random effect results from the reaction norm LMM estimating urinary cortisol concentrations (ng/ml SG) in response to time of day among immatures in five chimpanzee communities.*

| Groups | Name | Variance | Est.error | Q2.5 | Q97.5 |
|---|---|---|---|---|---|
| ID-year | Intercept | 0.30 | 0.08 | 0.14 | 0.46 |
| | Time of day | 0.27 | 0.08 | 0.10 | 0.41 |
| | Time of day$^2$ | 0.59 | 0.17 | 0.19 | 0.90 |
| Individual identity | Intercept | 0.25 | 0.09 | 0.06 | 0.41 |
| | Time of day | 0.12 | 0.08 | 0.01 | 0.30 |
| | Time of day$^2$ | 0.23 | 0.15 | 0.01 | 0.57 |
| Group-year | Intercept | 0.38 | 0.12 | 0.15 | 0.62 |
| Database code | Intercept | 0.44 | 0.17 | 0.20 | 0.84 |
| Residual | | 0.58 | 0.01 | 0.56 | 0.61 |

*Table S10: Medians of MCMC iterations (median) and their 5% and 95% quantiles for random effects included in the animal model and heritability calculations. Note that the technical predictor, database code, is excluded to compute the proportion of variance in inter-individual differences. The medians for all predictors combined are provided in Table S4.*

| **Predictor** | **Coefficient** | **Median proportion of variance** | **Lower CI** | **Upper CI** |
|---|---|---|---|---|
| Genetic | Intercept | 0.049 | 0.004 | 0.136 |
| | Time of day | 0.029 | 0.002 | 0.084 |
| | Time of $day^2$ | 0.055 | 0.005 | 0.162 |
| | Covariance (Intercept, Time of day) | 0.111 | -0.768 | 0.841 |
| | Covariance (Intercept, Time of $day^2$) | -0.119 | -0.842 | 0.764 |
| | Covariance (Time of day, Time of $day^2$) | -0.045 | -0.827 | 0.784 |
| Maternal identity | Intercept | 0.185 | 0.084 | 0.245 |
| | Time of day | 0.049 | 0.004 | 0.096 |
| | Time of $day^2$ | 0.053 | 0.005 | 0.156 |
| | Covariance (Intercept, Time of day) | 0.301 | 0.604 | 0.856 |
| | Covariance (Intercept, Time of $day^2$) | -0.206 | -0.847 | 0.720 |
| | Covariance (Time of day, Time of $day^2$) | -0.028 | -0.815 | 0.0795 |
| Individual identity | Intercept | 0.078 | 0.008 | 0.188 |
| | Time of day | 0.031 | 0.003 | 0.086 |
| | Time of $day^2$ | 0.061 | 0.006 | 0.174 |
| | Covariance (Intercept, Time of day) | 0.095 | -0.766 | 0.835 |
| | Covariance (Intercept, Time of $day^2$) | -0.118 | -0.834 | 0.757 |
| | Covariance (Time of day, Time of $day^2$) | -0.062 | -0.830 | 0.782 |
| ID-year | Intercept | 0.223 | 0.184 | 0.267 |
| | Time of day | 0.055 | 0.005 | 0.134 |
| | Time of $day^2$ | 0.179 | 0.022 | 0.337 |

| | | | | |
|---|---|---|---|---|
| | Covariance (Intercept, Time of day) | -0.137 | -0.763 | 0.604 |
| | Covariance (Intercept, Time of day$^2$) | -0.330 | -0.742 | 0.416 |
| | Covariance (Time of day, Time of day$^2$) | -0.146 | -0.842 | 0.740 |
| Group-year | Intercept | 0.583 | 0.452 | 0.747 |
| | Time of day | 0.080 | 0.011 | 0.173 |
| | Time of day$^2$ | 0.153 | 0.019 | 0.342 |
| | Covariance (Intercept, Time of day) | 0.105 | -0.672 | 0.748 |
| | Covariance (Intercept, Time of day$^2$) | -0.750 | -0.971 | 0.248 |
| | Covariance (Time of day, Time of day$^2$) | -0.121 | -0.807 | 0.717 |
| Database-code | Intercept | 0.584 | 0.306 | 1.071 |
| | Time of day | 0.210 | 0.106 | 0.384 |
| | Time of day$^2$ | 0.164 | 0.019 | 0.395 |
| | Covariance (Intercept, Time of day) | -0.668 | -0.924 | -0.037 |
| | Covariance (Intercept, Time of day$^2$) | 0.074 | -0.705 | 0.791 |
| | Covariance (Time of day, Time of day$^2$) | -0.094 | -0.773 | 0.693 |
| Residual error | | 0.650 | 0.639 | 0.660 |

*Table S11.Summary of genetic and maternal effect estimates on circadian cortisol responses in wild chimpanzees when group is included as predictor. Each coefficient represents a different component of the cortisol response. We also report the proportion of permutations for which these coefficient estimates were less than in the observed data. Coefficients in bold were larger in our observed data than in at least 95% of our random permutations.*

| Coefficient | Estimate | (lCI, uCI) | Proportion observed < permutations |
|---|---|---|---|
| *Genetic effect* | | | |
| $h^2_{intercept}$ | 0.007 | (0.000, 0.062) | 0.89 |
| $h^2_{linear}$ | 0.033 | (0.000, 0.421) | 0.75 |
| $h^2_{quadratic}$ | 0.041 | (0.000, 0.357) | 0.75 |
| *Maternal effect* | | | |
| **$m^2_{intercept}$** | **0.076** | **(0.017, 0.179)** | **0.00** |
| $m^2_{linear}$ | 0.082 | (0.001, 0.509) | 0.07 |
| $m^2_{quadratic}$ | 0.027 | (0.000, 0.266) | 0.38 |

*Table S12. Summary of genetic and maternal effect estimates on circadian cortisol responses in wild chimpanzees when only samples from Taï groups were used. Each coefficient represents a different component of the cortisol response. We also report the proportion of permutations for which these coefficient estimates were less than in the observed data. Coefficients in bold were larger in our observed data than in at least 95% of our random permutations.*

| Coefficient | Estimate | (lCI, uCI) | Proportion observed < permutations |
|---|---|---|---|
| *Genetic effect* | | | |
| $h^2_{intercept}$ | 0.009 | (0.000, 0.007) | 0.71 |
| $h^2_{linear}$ | 0.066 | (0.001, 0.279) | 0.60 |
| $h^2_{quadratic}$ | 0.027 | (0.000, 0.489) | 0.83 |
| *Maternal effect* | | | |
| **$m^2_{intercept}$** | **0.079** | **(0.001, 0.167)** | **0.02** |
| $m^2_{linear}$ | 0.128 | (0.001, 0.634) | 0.15 |
| $m^2_{quadratic}$ | 0.024 | (0.000, 0.238) | 0.20 |

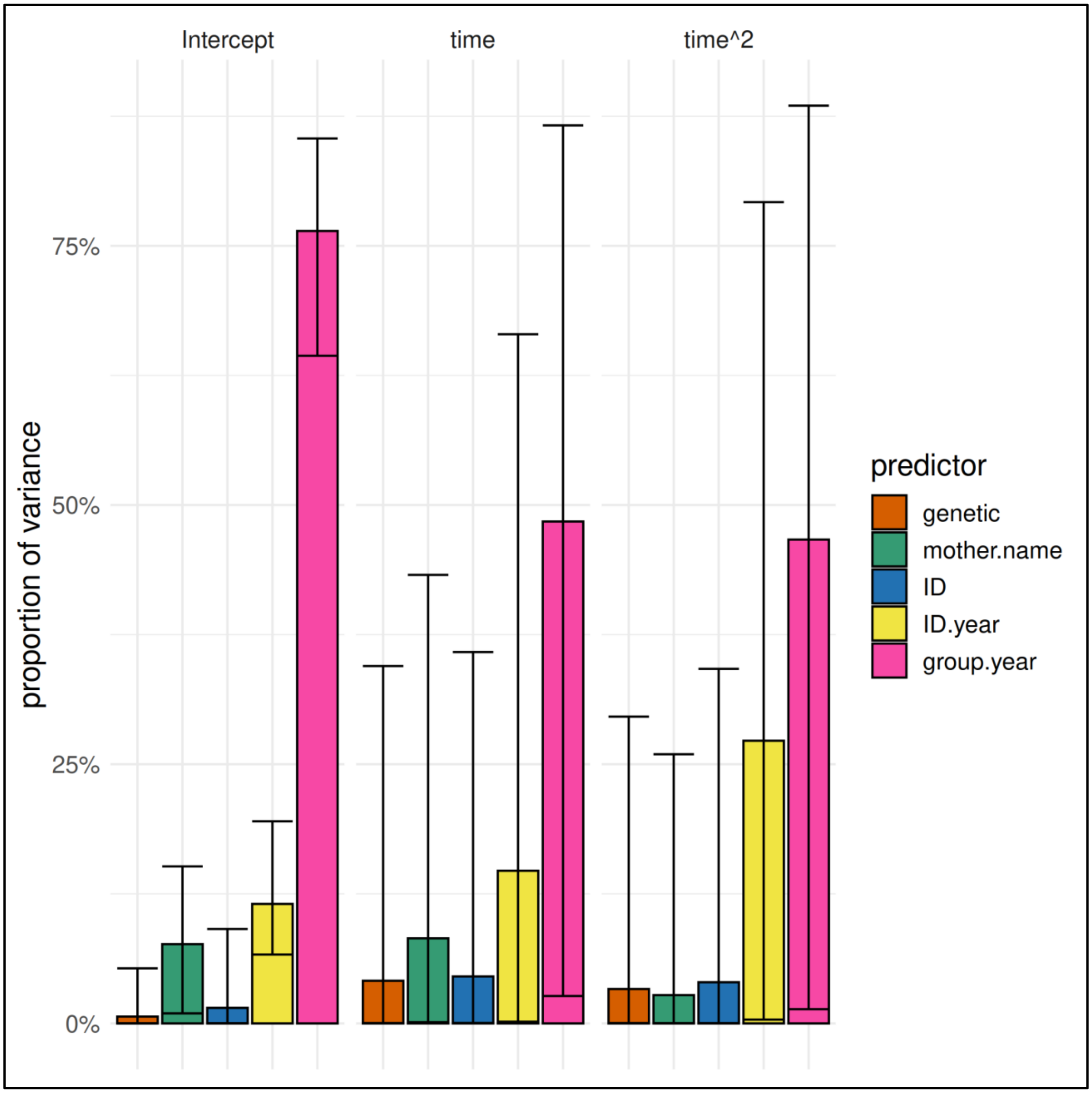


*Figure S6: Estimates for the proportion of variance in cortisol in wild chimpanzees explained by random effects for a model in which community is used as predictor instead of group size. The error bars represent the 95% credible interval range of the estimates.*

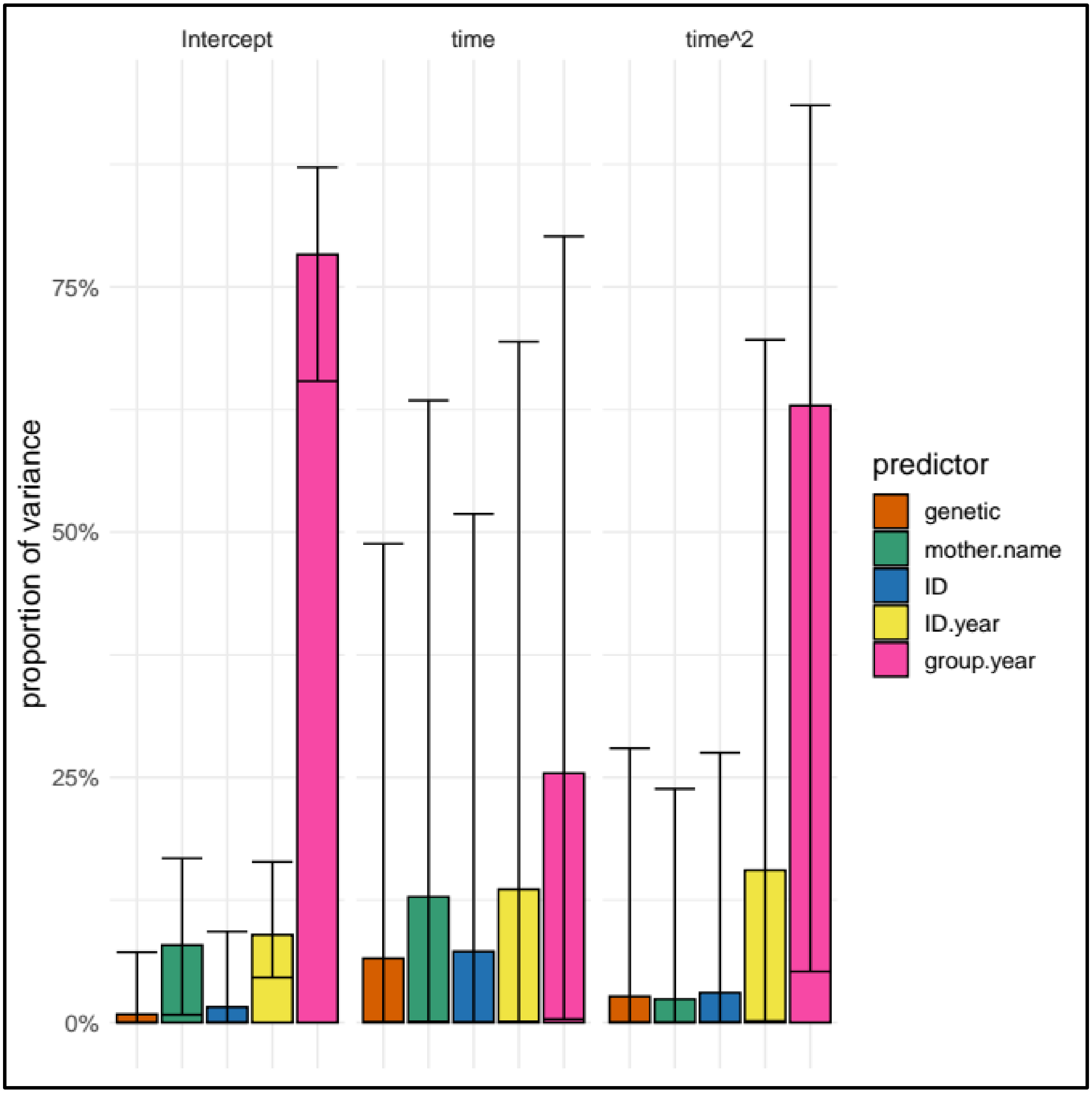


*Figure S7: Estimates for the proportion of variance in cortisol in wild chimpanzees explained by random effects for a model in which only samples from the Taï population are used. The error bars represent the 95% credible interval range of the estimates.*

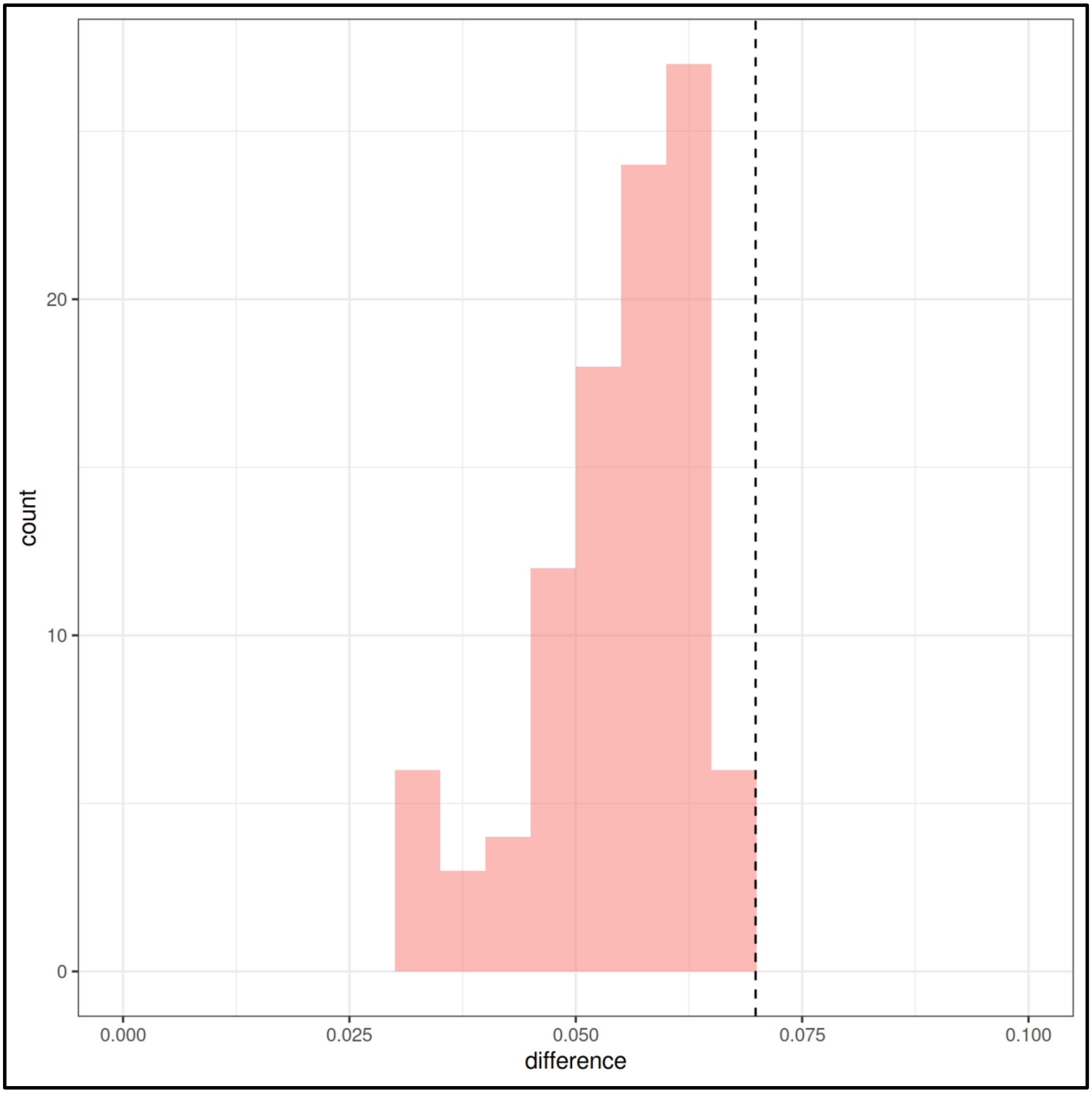


*Figure S8: Estimates of the difference in the proportion of variance explained by the maternal effect and that explained by genetic factors in the observed data (dashed line) and in 100 permutations of the data (red histogram), for the model in which community is used as a predictor rather than group size.*

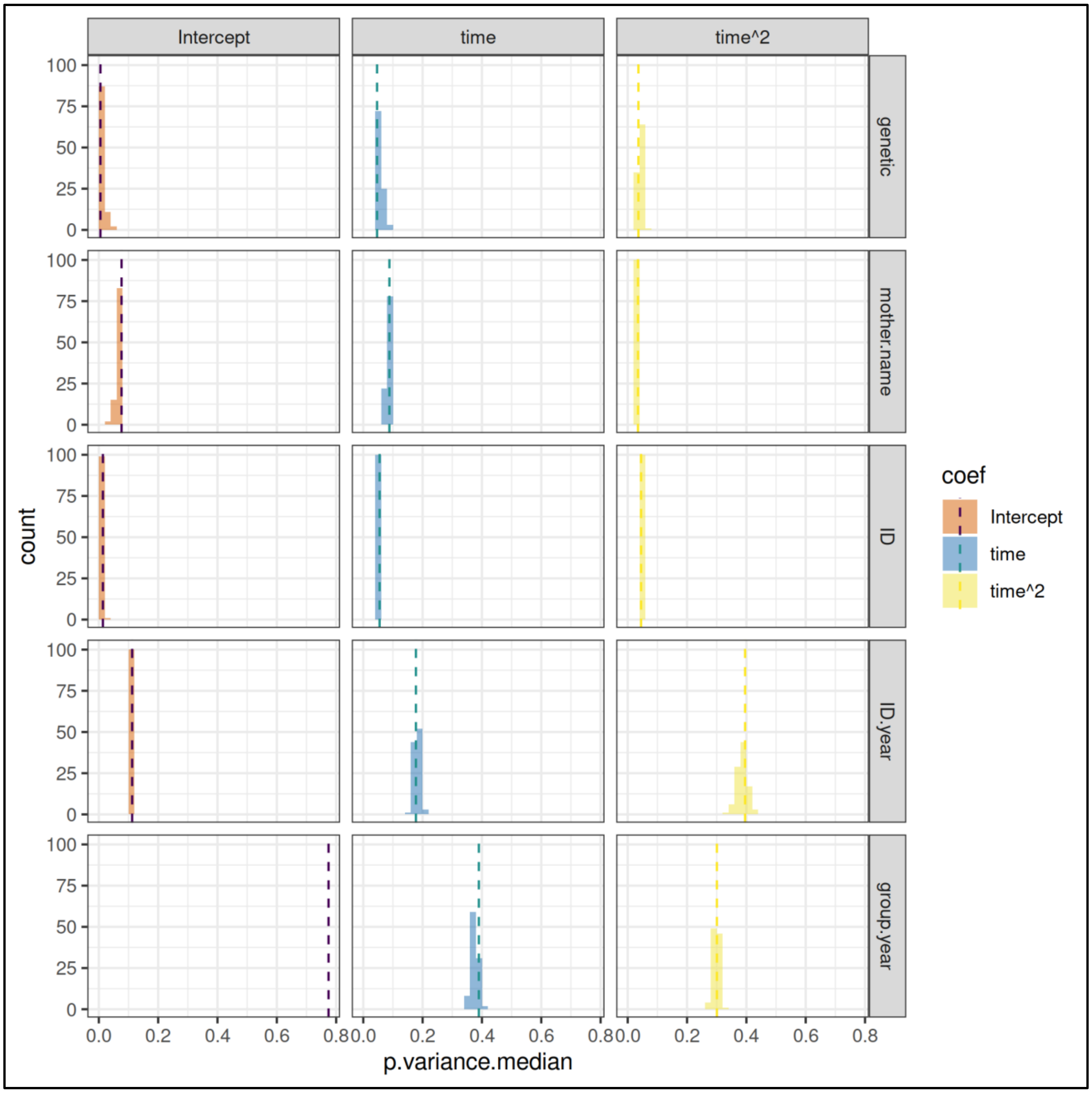


*Figure S9: Median proportion of variance estimates obtained from the observed data (dashed vertical lines) versus estimates obtained by 100 datasets with permuted genetic relationships between individuals. Histograms represent the counts of each estimate value from the permutations.*
